# Rules of engagement: determinants of chemokine receptor activation and selectivity by CCL27 and CCL28

**DOI:** 10.1101/2025.05.27.656466

**Authors:** Mian Huang, Aura F. Celniker, Rezvan Chitsazi, Douglas P. Dyer, Ariane L. Jansma, Irina Kufareva, Catherina L. Salanga, Tracy M. Handel

**Affiliations:** Skaggs School of Pharmacy and Pharmaceutical Sciences, University of California, San Diego, La Jolla, CA, 92093, USA; Graduate Program in Molecular Medicine, University of Maryland School of Medicine, Baltimore, MD, 21201 USA; MD Anderson Cancer Center, Houston, TX, 77030, USA; Wellcome Centre for Cell-Matrix Research, Manchester Academic Health Science Centre, University of Manchester, M13 9PT Manchester, UK; Geoffrey Jefferson Brain Research Centre, Manchester Academic Health Science Centre, University of Manchester, M6 8FJ Manchester, UK; Department of Chemistry, Point Loma Nazarene University, San Diego, CA, 92106, USA

**Keywords:** CCL27, CCL28, C-C chemokine receptor type 3 (CCR3), C-C chemokine receptor type 10 (CCR10), cell migration, glycosaminoglycan, superagonist

## Abstract

The distinct functional roles of chemokines CCL27 and CCL28 in epithelial immunity of skin and mucosal tissues, respectively, are coordinated by their shared receptor, CCR10 and the CCL28-specific receptor, CCR3. In this study, we conducted structure-function studies focused on the N-termini of these two chemokines to identify determinants of receptor activation, internalization and binding specificity. Deletion of two N-terminal residues of CCL27 resulted in a CCR10 antagonist, highlighting the critical roles of these residues in driving receptor pharmacology. Extension with a Phe produced a superagonist by occupying an available subpocket in the receptor binding site. Swapping the N-terminus of CCL28 onto the CCL27 globular domain (NT28-CCL27) also resulted in a superagonist of CCR10, but the opposite swap (NT27-CCL28) showed equivalent or reduced activity compared to WT CCL28, indicating that the CCL28 N-terminus is a stronger driver of CCR10 signaling. The effect of these and other mutations were rationalized by AlphaFold models of the CCR10 complexes. AlphaFold modeling also revealed that the reduced size of the binding pocket, and more basic nature of the N-terminus and extracellular loops of CCR3 compared to CCR10, contribute to its specificity for CCL28 while CCR10 accommodates both ligands. The overall basic nature of CCL28 also contributes to its high affinity for glycosaminoglycans, which is likely important for its retention in mucosal tissues. These data illustrate the modular compositions of these chemokines that has evolved to achieve overlapping but non-redundant functions, and the exploitation of this modular nature to produce engineered chemokines for probing or targeting CCR10 in disease contexts.

## Introduction

Chemokines and their receptors belong to a large family of chemoattractant proteins that control cell movement and positioning during development and homeostatic processes, as well as during inflammatory responses to infection and other physiological insults (1, 2). As central mediators of inflammation, they have been pursued as potential therapeutic targets in numerous inflammatory, autoimmune and infectious diseases (3–5). CC chemokines, CCL27 and CCL28, both ligands of CC chemokine receptor 10 (CCR10), are critical regulators of epithelial immunity, but play functionally unique roles as suggested by their distinct expression patterns. CCL27 is constitutively expressed by skin keratinocytes (6) and CCL28 by epithelial cells of various mucosal tissues including GI tract, salivary glands, mammary glands, nasal epithelial cells and lung (7–9). CCR10 is expressed on skin homing T cells, including cutaneous lymphocyte-associated antigen positive memory T cells (6, 10, 11), T regulatory cells (Tregs) (12), as well as melanocytes, dermal fibroblasts, and dermal microvascular endothelial cells (13) that respond to constitutively produced or upregulated CCL27. CCR10 is also expressed by mucosal IgA antibody-producing plasma blasts and plasma cells that respond to CCL28 (14). The potential of CCR10 as a therapeutic target is suggested by its role in T cell-mediated skin inflammation such as psoriasis (6) and atopic or allergic-contact dermatitis (10, 15), in addition to its expression on lung epithelial cells in humanized mouse models of idiopathic pulmonary fibrosis (IPF) (16). Additionally, CCR10 contributes to various cancers both through its expression directly on malignant cells and by its expression on cells in the tumor microenvironment (TME). For example, engagement of CCL27 by CCR10 expressed on skin melanoma cells promotes immune evasion (17) whereas its expression on Tregs and the CCL28-mediated recruitment of those cells into the TME of ovarian and liver cancer promotes tumor growth through the creation of an immunosuppressive microenvironment (18, 19). Thus, blocking CCR10 may be a useful approach for skin inflammatory diseases, IPF and enhancing the efficacy of cancer therapies (20, 21).

In addition to CCR10, CCL28 also binds CC chemokine receptor 3 (CCR3), which is expressed on CD4+ T cells in nasal mucosa (22), as well as eosinophils (8, 23), basophils, and neutrophils (24) following infection. CCR3 is a promiscuous receptor with numerous chemokine ligands (e.g., CCL5, CCL7, CCL11, CCL13, CCL15, CCL24, CCL26 and CCL28 (25)), although most exhibit little to no expression on normal nasal mucosa (22), suggesting a specific role of CCL28 and CCR3 homing of CD4+ T cells to the upper airways. CCR3 is implicated in numerous allergic diseases including asthma, eosinophilic esophagitis and atopic dermatitis (26, 27) making the CCR3 axis another attractive pharmaceutical target.

Given their importance in both normal physiology and disease, surprisingly little is known about the structure-function details of CCL27 and CCL28, apart from NMR structures (28, 29). Thus, to understand the molecular interactions of CCL27 and CCL28 with their receptors, we undertook a mutagenesis study focusing on the chemokine N-terminal residues, which are key determinants of receptor pharmacology. Studies of effects on signaling and internalization revealed variants of CCL27 that are superagonists of CCR10. AlphaFold models of WT and mutant chemokines in complex with CCR10 and CCR3 rationalized the functional consequences of the mutations, as well as the specificity of CCL27 for CCR10 and CCL28 for CCR10 and CCR3. Our functional data and computational models also provide a starting point for further optimization of CCL27 superagonists that may have utility as adjuvants for vaccine development (30, 31). In addition to signaling receptors, chemokines interact with glycosaminoglycans (GAGs) (32, 33), which contributes to their localization on cell surfaces and the extracellular matrix (34). Here we demonstrate that CCL27 and CCL28 exhibit markedly different GAG-binding propensities that correlate with their accumulation on cells. Along with their distinct expression profiles, these findings underscore the non-redundant roles of the two chemokines, despite sharing CCR10 as a common receptor.

## Results

### N-terminal mutations of CCL27 produce CCR10 antagonists

All chemokines share a globally similar architecture when engaging their receptors, whereby the chemokine globular domain rests on the extracellular face of the receptor and interacts with the receptor N-terminus (regions referred to as chemokine recognition sites 1 and 0.5, CRS1 and CRS0.5) and extracellular loops (ECLs), while the chemokine N-terminus reaches into the transmembrane receptor binding pocket (CRS2) and promotes activation through rearrangements of the transmembrane helices (35–37). Accordingly, the N-termini of most chemokines drive the pharmacological responses of their receptors (38–41). Extension of some chemokines by even a single amino acid can profoundly increase or decrease efficacy (41–43). Truncation of chemokine N-termini can be activating or deactivating with effects on receptor binding affinity, signaling and trafficking (41, 44), and proteolytic N-terminal processing represents a natural mechanism for regulating chemokine activity (45–47). We therefore sought to characterize the effect of N-terminal modifications of CCL27 and CCL28 on receptor signaling and internalization and how it relates to their receptor specificity (**Fig. 1A-D**).

**Figure 1.**
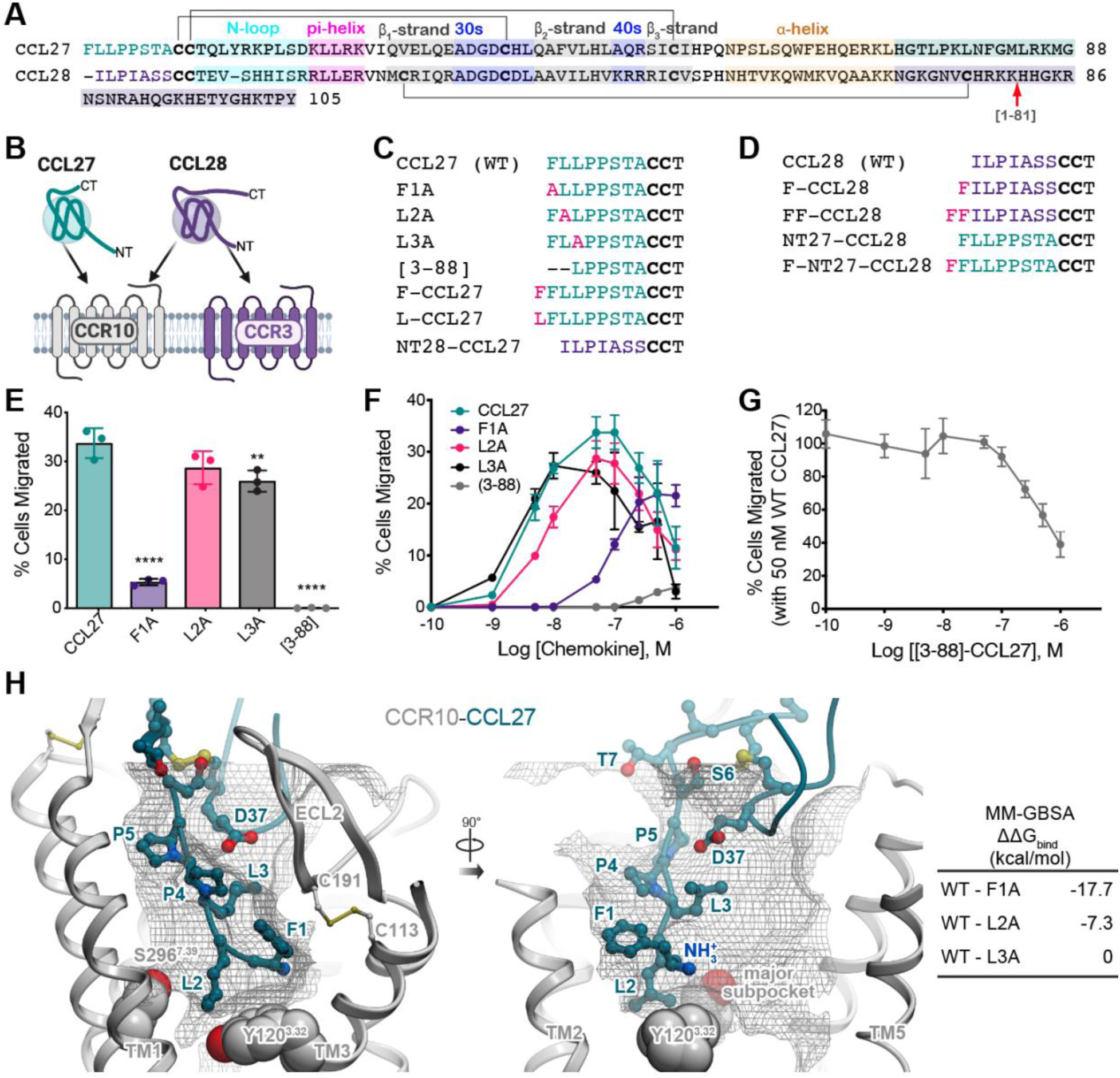
Identification of N-terminally modified CCL27 mutants that function as CCR10 antagonists. (**A**) A sequence alignment of CCL27 and CCL28 where the N-terminal residues are colored teal and purple, respectively, and characteristic chemokine features labeled accordingly. Disulfide bonds are indicated by black lines, the CCL28 C-terminal truncation mutant is indicated by a red arrow (residue 81 of CCL28) and the extended C-terminal domains are highlighted. (**B**) Cartoon depicting CCL27 specificity for CCR10 and CCL28 for CCR3 and CCR10. (**C**) Sequences of WT and N-terminally modified mutants of CCL27 and (**D**) CCL28. Disulfide bonded Cys are bolded. (**E**) Cell migration analysis of WT CCL27 and N-terminal mutants with CCR10-expressing L1.2 cells at 50 nM (average concentration for maximal migration of WT CCL27) and shown as representative data (mean ± SD) of three independent experiments performed in technical triplicates. Statistical analysis was performed by ordinary one-way ANOVA with *post hoc* Dunnett’s multiple comparisons test (∗∗∗∗p< 0.0001; ∗∗p< 0.01; CCL27 : L2A, p = 0.0673). (**F**) migration across a broad chemokine concentration range (0.1 nM to 1 μM), plotted as the percent of cells migrated relative to maximal migration (i.e., no filter). Shown are representative data (mean ± SD) of three independent experiments performed in technical triplicates. (**G**) Competition chemotaxis binding assay of CCL27 with the N-terminal truncation mutant [3-88]. WT CCL27 (50 nM) was incubated with increasing amounts of [3-88]-CCL27 and the percent cell migration was compared after a 2 h incubation period at 37°C. Shown are representative data (mean ± SD) of three independent experiments performed in technical triplicates. (**H**) Structural model of the N-terminus and 30s loop of WT CCL27 (teal sticks) in complex with CCR10 (gray ribbon). The optimal ligand atom placement surface in the orthosteric pocket of the receptor is shown as a gray mesh. Some TM helices of the receptor are hidden for clarity. Receptor residues Tyr120(3.32) and Ser296(7.39) are shown in gray spheres. Predicted Prime MM-GBSA energy difference values for the N-terminal mutants (F1A, L2 and L3A) of CCL27 relative to WT CCL27.

As a starting point, three N-terminal Ala mutants of CCL27 (F1A-, L2A- and L3A-CCL27) and an N-terminal truncation mutant ([3-88]-CCL27) (**Fig. 1C**) were generated and characterized in several functional assays. In transwell migration assays with murine L1.2 pre-B cells stably expressing human CCR10, 50 nM L2A and L3A showed ∼15-20% reduction in migration compared to WT CCL27, while F1A and [3-88]-CCL27 practically abrogated migration (a reduction of least 85%, **Fig. 1E**). Full dose response curves revealed an approximately 35% reduction in efficacy and 10-fold reduction in potency for F1A-CCL27 relative to WT CCL27. The truncation mutant [3-88]-CCL27 was unable to induce cell migration up to 1 μM chemokine (**Fig. 1F**), despite preserved receptor binding and despite being able to block cell migration to WT CCL27 (**Fig. 1G**).

To understand the structural basis for the role of the first three residues of CCL27 in receptor activation, an AlphaFold model of CCL27 bound to CCR10 was generated (**Fig. 1H**). In the model, residues Phe1 and Leu2 of the chemokine directly interact with Tyr120(3.32) and Ser296(7.39) at the bottom of the orthosteric binding pocket of the receptor (48); these pocket residues are known to be involved in the activation of multiple homologous receptors (49–53). Phe1 of CCL27 also makes favorable hydrophobic packing interactions under the extracellular loop 2 (ECL2) and the conserved disulfide bridge Cys113(3.25)-Cys191(ECL2) of the receptor (**Fig. 1H**). Leu3 may also provide hydrophobic interactions with the pocket due to its proximity to CCR10 L193(ECL2). The Prime molecular mechanics generalized Born and surface area solvation (MM-GBSA) calculation ((54, 55), Schrödinger, LLC, New York, NY, 2021) predicted an energy difference of 17.7 kcal/mol between WT CCL27 and F1A, compared to 7.3 kcal/mol for L2A and no difference for L3A, consistent with the impact of these mutations on CCL27-induced migration. Together these findings demonstrate that a primary determinant of receptor activation resides within the first two residues of CCL27 and that [3-88]-CCL27 is a CCR10 antagonist.

### A phenylalanine addition to the N-terminus of CCL27 produces a CCR10 superagonist

We next investigated the impact of adding residues to the N-termini of CCL27 and CCL28 since such additions can also modulate signaling, sometimes resulting in useful variants with superagonist or antagonist properties (41–43, 56) and potential therapeutic value (41–43). Because these modifications often involve bulky groups, we extended CCL27 with an N-terminal Phe to produce F-CCL27 (**Fig. 1C**) and observed markedly enhanced agonist activity in multiple functional assays (**Fig. 2**). In a bare filter chemotaxis assay involving CCR10-expressing L1.2 cells, the efficacy and potency of F-CCL27 were 1.1- and 10-fold higher, respectively, than those of WT CCL27 (**Fig. 2A**). F-CCL27 also exhibited a 2.7-fold higher efficacy and 3-fold greater potency than WT CCL27 in promoting transendothelial cell migration (**Fig. 2B**); the difference in efficacy was particularly striking compared to the bare filter migration assay (**Fig. 2A** vs. **2B**).

**Figure 2.**
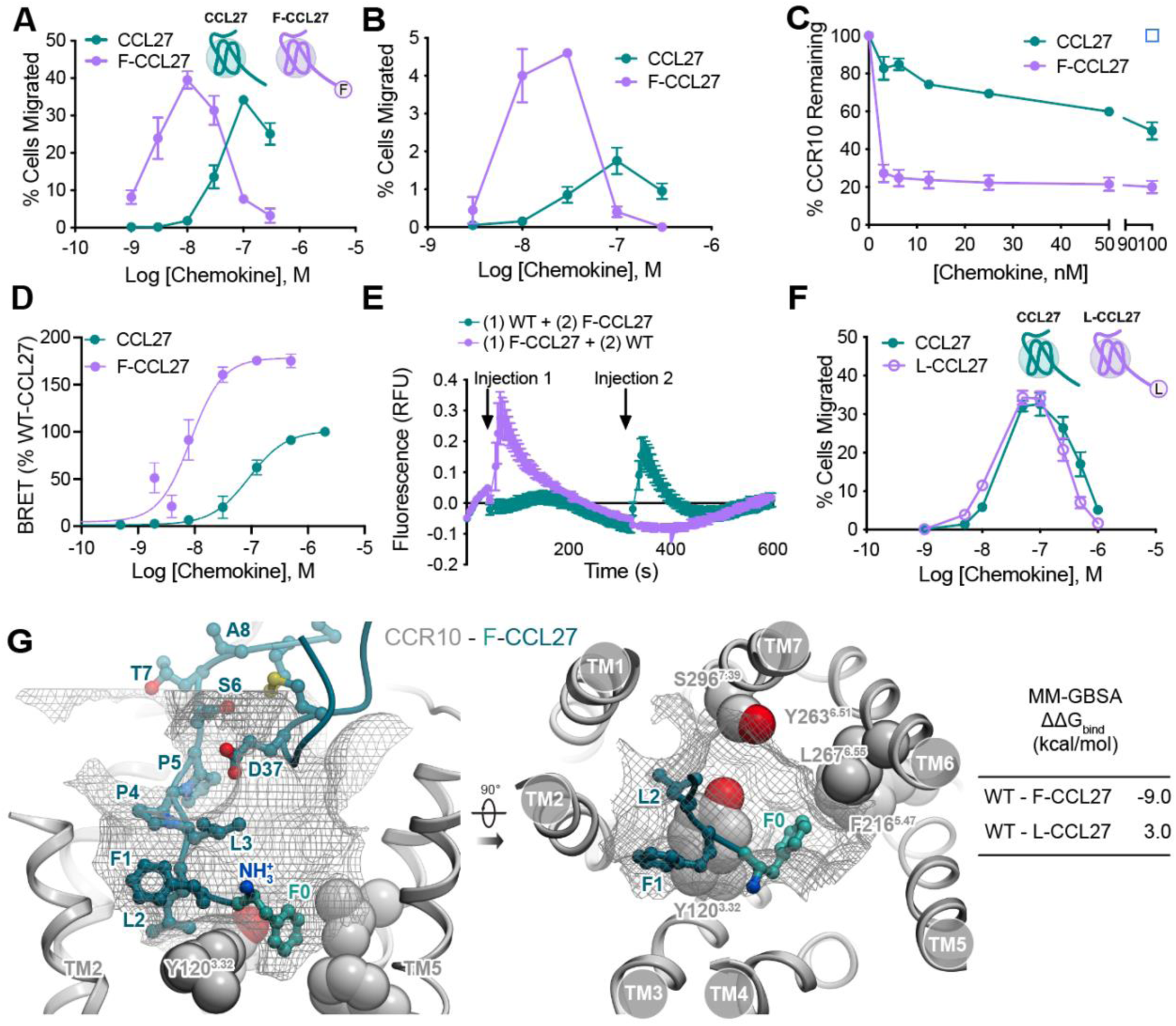
Functional analysis of the CCR10 superagonist, F-CCL27. (**A**) Cell migration analysis of CCL27 WT and F-CCL27 with L1.2/CCR10 cells across varying chemokine concentrations. Results shown are representative data (mean ± SD) of independent experiments (n=6) performed in technical duplicates or triplicates. (**B**) Transendothelial cell migration of WT CCL27 and F-CCL27 with L1.2/CCR10 cells across varying chemokine concentrations with data plotted as the percent of cells migrated after a 2 h incubation at 37°C. Shown are representative data (mean ± SD) of independent experiments (n=3) performed in technical duplicates. (**C**) WT or F-CCL27-mediated internalization of L1.2/CCR10 cells. Cells were treated with varying concentrations of chemokine, incubated at 37°C for 45 min and then receptor levels were detected and plotted as the percent of receptor remaining compared to cells treated with no chemokine. Results shown are the mean ± SEM of single measurements obtained for each concentration combined from at least three independent experiments (n=3-4). (**D**) β-arrestin recruitment by BRET. Cells were transiently transfected with CCR10-RlucII and βarr2-GFP10 and then stimulated with varying concentrations of chemokine. The BRET ratios are shown as a % of WT-CCL27. Results shown are the mean ± SD of independent experiments (n=4) performed in technical duplicates or triplicates. (**E**) Chemokine-mediated desensitization of L1.2/CCR10 cells as measured by calcium flux. Cells were treated with an initial dose of 20 nM WT or F-CCL27 (injection 1) followed by a second addition of 20 nM F-CCL27 and WT CCL27 (injection 2), respectively, after the 75th reading cycle (at approximately 320 s). Shown are representative data (mean ± SD) of independently performed experiments (n=3) performed in technical triplicates. (**F**) Cell migration analysis of CCL27 WT and L-CCL27 with L1.2/CCR10 cells across varying chemokine concentrations plotted as the percent of cells migrated after a 2 h incubation at 37°C. Shown are representative data (mean ± SD) of independent experiments (n=3) performed in technical triplicates. (**G**) Structural model of the N-terminus and the 30s loop of F-CCL27 (teal sticks) in complex with CCR10 (gray ribbon). The optimal ligand atom placement surface in the orthosteric pocket of the receptor is shown as gray mesh. Some TM helices of the receptor are hidden for clarity. Receptor residues Tyr120(3.32), F216(5.47), Y263(6.51), L267(6.55), and Ser296(7.39) are shown as gray spheres. Predicted Prime MM-GBSA energy difference values for the N-terminal mutants F-CCL27 and L-CCL27 relative to WT CCL27.

In addition to its effect on G protein-mediated signaling responses, we also investigated whether the addition of the N-terminal Phe impacted the ability of the receptor to internalize following chemokine stimulation using flow cytometry **(Fig. 2C**). F-CCL27 internalized CCR10 much more effectively than WT CCL27, with only ∼30% of the receptor remaining on the cell surface after 45 min compared to ∼80% after stimulation with WT CCL27 at the lowest concentration tested (3.1 nM) (**Fig. 2C**). Since internalization is often a β-arrestin mediated process (57), we also used a BRET-based association assay to monitor β-arrestin recruitment to the receptor (58) with CCR10-RlucII and GFP10-tagged β-arrestin2 (βarr2-GFP10) as BRET donor-acceptor pairs (**Fig. 2D**). Compared to WT-CCL27, which promoted β-arrestin recruitment to CCR10 with an EC_50_ of 71 nM, F-CCL27 recruited β-arrestin with an EC_50_ of 6.8 nM (**Fig. 2D**). Further characterization of 20 nM F-CCL27 in a calcium flux assay showed a robust response following the first injection of chemokine compared to the equivalent concentration of WT CCL27, which exhibited little to no signal (**Fig. 2E, Supp. Fig. 1**). Notably, F-CCL27 fully desensitized CCR10 as shown by the lack of a response to subsequent stimulation by WT CCL27, whereas cells initially stimulated with WT CCL27 produced a robust secondary response to F-CCL27, almost as large as the initial response with F-CCL27 (**Fig. 2E**). By contrast with Phe0, the addition of a N-terminal Leu0 (L-CCL27) showed little to no change compared to WT in a bare filter chemotaxis assay (**Fig. 2F**), indicating the specificity of the Phe addition for enhanced activation of CCR10. Taken together, the more potent responses of F-CCL27 with respect to receptor activation, desensitization and internalization, suggest that it is a superagonist. On the other hand, analogous N-terminal modifications in CCL28 (F-CCL28 and FF-CCL28, **Fig. 1D**) had little to no effect on CCR10-mediated migration compared to the WT chemokine in a bare filter chemotaxis assay (**Supp. Fig. 2**), suggesting marked differences in how these ligands activate CCR10.

To investigate the structural mechanism underlying F-CCL27 superagonism, we generated an AlphaFold model of CCR10 bound to F-CCL27 (**Fig. 2G**). F-CCL27 shares most of WT CCL27 interactions with CCR10; however, the N-terminal Phe of F-CCL27 extends into the major subpocket of CCR10 and directly contacts multiple residues in the receptor TM helices 5 and 6. Specifically, Phe0 makes direct contacts with Leu267(6.55) and, in some of the predicted conformations, with Tyr263(6.51), both part of a hydrophobic stack in TM6 just above the “toggle switch” residue 6.48 (which in CCR10 is a Gln, in contrast to the majority of GPCRs that have a Trp at the corresponding position). These contacts, absent in WT CCL27, likely explain the superagonist properties of F-CCL27. Consistent with experimental observations, the Prime MM-GBSA energy difference with WT CCL27 was highly favorable for the superagonist F-CCL27 (9 kcal/mol, **Fig. 2G**), whereas L-CCL27 was less favorable by 3 kcal/mol.

### Fusing the N-terminus of CCL28 onto the core domain of CCL27 enhances CCR10 activation

To further understand the contributions of the N-termini of CCL27 and CCL28 to receptor specificity and engagement, we generated N-terminal chimeras (**Fig. 1C, D**) and examined their ability to activate CCR10 (**Fig. 3**). NT28-CCL27, a chimera consisting of the N-terminus of CCL28 and the globular core domain of CCL27, showed increased potency and efficacy in promoting the migration of L1.2/CCR10 cells compared to WT CCL27 (**Fig. 3B**), while NT27-CCL28 and F-NT27-CCL28 were equipotent to WT CCL28 (**Fig. 3C, Supp. Fig. 2**). CCL28 was also more potent than CCL27 in inducing internalization of CCR10 and the addition of the CCL28 N-terminus onto CCL27 conferred NT28-CCL27 with a stronger capacity to internalize CCR10 than WT CCL27 (**Fig. 3D**). By contrast, NT27-CCL28 was less potent in internalizing CCR10 than WT CCL28, consistent with the more modest internalizing capacity of CCL27 (**Fig. 3D**). NT28-CCL27 also promoted β-arrestin recruitment to CCR10 with a 1.5-fold greater efficacy and 3.5-fold greater potency compared to WT CCL27 (**Supp. Fig. 3**). Thus, despite the relatively low sequence homology between the chemokine globular domains (∼30% identity), they still bind CCR10 in a manner that accommodates exchange of the chemokine N-termini with preservation of some of the activation characteristics (e.g., internalization) of the chemokine from which the N-terminus was derived.

**Figure 3.**
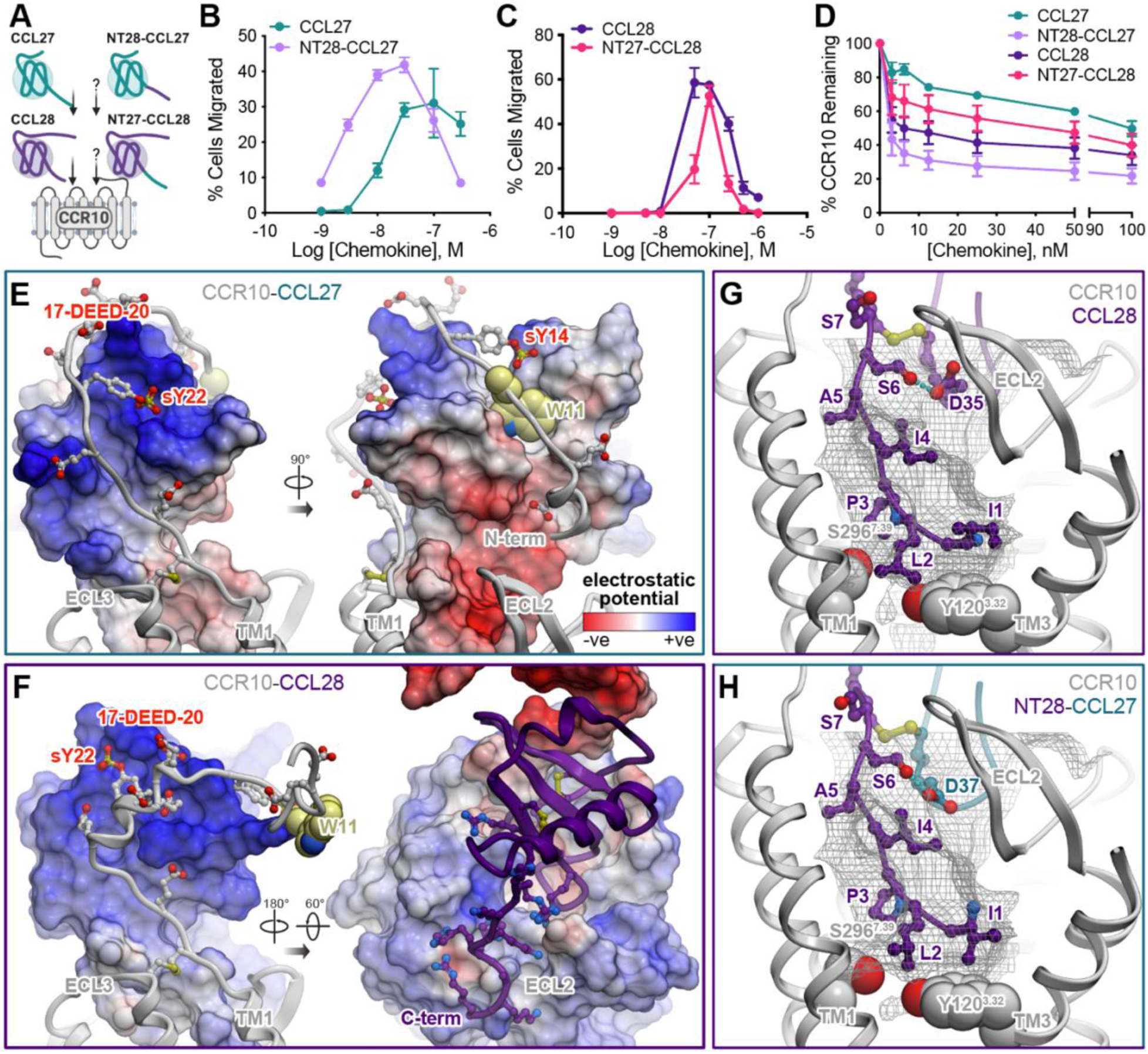
Pharmacological and structural characterization of N-terminally swapped chimeras of CCL27 and CCL28. (**A**) Cartoon depicting CCL27 and CCL28 specificity for CCR10 and unknown activity of N-terminal chimeras. (**B, C**) Cell migration analysis with L1.2/CCR10 cells across varying chemokine concentrations of (**B**) WT CCL27 and NT28-CCL27. Shown are representative data (mean ± SD) of independent experiments (n=5) performed in technical duplicates or triplicates and plotted as the percent of cells migrated after a 2 h incubation at 37°C. (**C**) WT CCL28 and NT27-CCL28. Shown are representative data (mean ± SD) of independent experiments (n=3) performed in technical triplicates and plotted as the percent of cells migrated after a 2 h incubation at 37°C. (**D**) Comparison of WT or mutant chemokine-mediated internalization of L1.2/CCR10 cells. Cells were treated with varying concentrations of chemokine, incubated at 37°C for 45 min and then receptor levels were detected and plotted as the percent of receptor remaining compared to cells treated with no chemokine. Results shown are the mean ± SEM of single measurements obtained for each concentration combined from at least three independent experiments (n=3-4). (**E**) Interactions of the receptor N-terminus with the globular core in the chemokine in the structural model of the CCR10-CCL27 complex. The complex is shown in two orientations; the chemokine is represented as a surface mesh colored by the electrostatic potential; the receptor is shown as a ribbon and sticks. Receptor residue W11 is shown in light-yellow spheres. (**F**) Interactions of the receptor N-terminus with the globular core in the chemokine in the structural model of the CCR10-CCL28 complex. The complex is shown in two orientations. On the left, chemokine is represented as a surface mesh colored by the electrostatic potential while the receptor is shown as ribbon and sticks. On the right, the receptor is shown as the electrostatic potential mesh while the chemokine is in purple ribbon. **(G, H**) Structural models of CCR10 (white ribbon and spheres bound to CCL28 (**G**) and NT28-CCL27 (**H**). Chemokine segments originated from CCL28 are shown in purple sticks, from CCL27 in teal sticks. The optimal ligand atom placement surface in the orthosteric pocket of the receptor is shown as gray mesh. Some TM helices of the receptor are hidden for clarity. Residues Tyr120(3.32) and Ser296(7.39) of the receptor are shown in gray spheres. An intramolecular hydrogen bond between residues Ser6 and Asp35 of CCL28 (**G**) or Asp37 of CCL27 (**H**) is shown as a cyan dotted line.

To rationalize the experimental data, we generated structural models of CCR10 complexes with CCL27 and CCL28 using AlphaFold (**Fig. 1H, 3E-G**). The most striking differences between CCR10 recognition of CCL28 and CCL27 are observed in CRS1 (**Fig. 3E, F**). With CCL28, the proximal N-terminus of CCR10 including its acidic cluster 17-DEED-20 and the predicted sulfotyrosine sTyr22, folds into a short α-helix and favorably engages the chemokine’s basic 40s loop (46-KRR-48), whereas the distal N-terminus of CCR10 is largely unutilized (**Fig. 3F**, right). By contrast, when bound to CCL27, the N-terminus of CCR10 not only favorably interacts with the basic residues in its N-loop (Lys16), pi-helix (20-KLLRK-24), and 40s loop (R50), but also wraps around the chemokine (**Fig. 3E)**. On the opposite side of CCL27, Trp11 in the receptor N-terminus favorably packs in the pocket formed by the β_1_-strand and the unique loop between the two C-terminal helices of CCL27 (**Fig. 3E**, right); this pocket does not exist in CCL28 because it lacks the 2^nd^ C-terminal helix. These observations suggest that the globular core of CCL27 provides a more favorable context for CCR10 CRS1 binding, compared to the globular core of CCL28.

In CRS2, the N-terminus of CCL28 binds CCR10 in a manner similar to that of WT CCL27, with CCL28 Ile1, Leu2 and Ile4 occupying approximately the same areas and cavities in the binding pocket as CCL27 Phe1, Leu2 and Leu3, respectively (**Fig. 1H, 3G**). However, closer examination reveals differences in the conformation and interaction of the chemokines’ proximal N-termini. The shorter, 7 amino acid N-terminus of CCL28 directly descends into the binding pocket whereas the proximal part of the longer, 8 amino acid N-terminus of CCL27 forms a short antiparallel β-sheet with CCR10 residues 31-CYK-33 (**Supp. Fig. 4**). This, and the proximal N-terminal sequence of CCL28, enables a favorable intramolecular hydrogen bond in the chemokine, between Ser6 (Cys8 minus 2) and Asp35 (Cys36 minus 1) (**Fig. 3G**), potentially stabilizing the observed conformation. On the other hand, an analogous hydrogen bond in the predicted binding pose of CCL27 appears impossible because the residues in proximity of the corresponding Asp37 (C38 minus 1) of CCL27 are 4-PP-5, which lack hydrogen bond donor functionality (**Fig. 1H**). This suggests that the N-terminus of CCL28 interacts with the CRS2 of CCR10 more favorably than the N-terminus of CCL27.

When the N-terminus of CCL28 is grafted onto the globular core of CCL27 (NT28-CCL27), the result combines the best of both complexes including the favorable CRS1 interactions of the CCL27 globular core (**Fig. 3E**), CRS2 interactions of the CCL28 N-terminus (**Fig. 3G, H**) and the hydrogen bond between the proximal N-terminus and the 30s loop (**Fig. 3H**). This possibly explains the improved agonist properties of this chimera compared to both WT CCL27 and CCL28 (**Fig 3B, D**). By contrast, grafting the N-terminus of CCL27 onto the globular core of CCL28 (NT27-CCL28) does not produce such synergy, explaining the lack of agonist potency enhancement (**Fig. 3C**).

### Swapping the N-termini of CCL27 and CCL28 does not lead to CCR3 activation

In addition to CCR10, CCL28 activates CCR3, while CCL27 is a CCR10-selective ligand (**Fig. 4A**). In order to determine the requirements for CCR3 binding and activation and the specificity of the two chemokines, N-terminal chimeras of CCL27 and CCL28 were compared for their ability to promote migration of CCR3-expressing L1.2 cells in a barefilter chemotaxis assay. As shown in **Fig. 4B**, NT28-CCL27 was unable to promote migration or compete with another CCR3 ligand (CCL7-HA) for binding the receptor (**Fig. 4C**), suggesting this chimera has little or no affinity for CCR3. The NT27-CCL28 chimera was also incapable of activating CCR3 (**Fig. 4B**) even though it still competed with CCR3 agonist CCL7 for receptor binding (**Fig. 4C**). This suggests that the N-terminus and the globular domain of CCL28 work together to determine receptor selectivity and agonist activity, and that the N-terminal domain of CCL27 is incompatible with CCR3.

**Figure 4.**
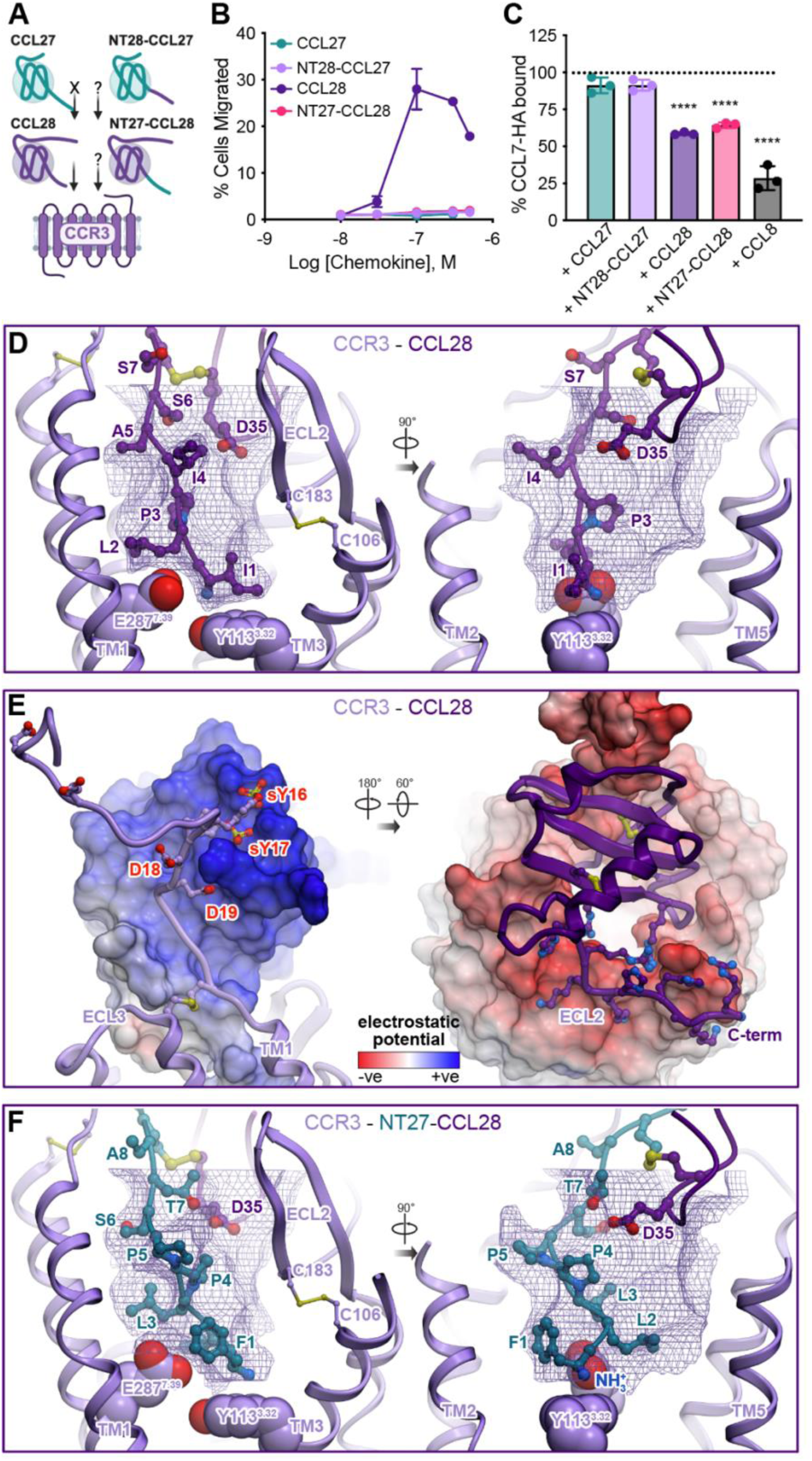
Functional and structural analysis of N-terminal chimeras of CCL27 and CCL28 with CCR3. (**A**) Cartoon depicting CCL28 specificity for CCR3, lack of activity by CCL27 and unknown activity of N-terminal chimeras. (**B**) Cell migration analysis of WT CCL27, WT CCL28 and N-terminal chimera mutants (NT28-CCL27 and NT27-CCL28) with L1.2/CCR3 cells across varying chemokine concentrations. Shown are representative data (mean ± SD) of independent experiments (n=3) performed in technical duplicates, plotted as the percent of cells migrated after 2 h incubation at 37°C. (**C**) Competition binding experiments were performed on L1.2/CCR3 cells using C-terminally HA-tagged CCL7 (CCL7-HA). Cells were incubated with 250 nM of CCL7-HA alone or in combination with a 4-fold excess of competing ligand (1000 nM) for 30 min on ice, cells were washed and then CCL7 levels were determined by flow cytometry analysis using an anti-HA antibody. Results shown are the mean ± SEM of single measurements obtained for each concentration combined from independent experiments (n=3); the dotted line depicts 100% retention of bound CCL7-HA. Statistical analysis was performed by ordinary one-way ANOVA with *post hoc* Dunnett’s multiple comparisons test. (∗∗∗∗p< 0.0001; CCL7-HA : + CCL27, p = 0.1026; CCL7-HA : + NT28-CCL27, p = 0.1026). (**D-F**) Structural model of CCR3 (violet ribbon and spheres) bound to CCL28 (**D, E**) or NT27-CCL28 (**F**). Chemokine segments originated from CCL28 are shown in purple sticks and ribbons, from CCL27 in teal sticks and ribbons. (**D**) and (**F**) depict the predicted interactions of chemokine N-termini and 30s loops (sticks) with the orthosteric pocket of the receptor. The optimal ligand atom placement surface is shown as violet mesh. Some TM helices of the receptor are hidden for clarity. Residues Tyr113(3.32) and Glu287(7.39) of the receptor are shown in spheres. (**E**) depicts the interactions between the globular domain of CCL28 and the N-terminus (left) or extracellular loops (right) of the CCR3. The complex is shown in two orientations. On the left, chemokine is represented as a surface mesh colored by the electrostatic potential while the receptor is shown as ribbon and sticks. On the right, the receptor is shown as the electrostatic potential mesh while the chemokine is in purple ribbon.

To further dissect the molecular details underlying chemokine selectivity for CCR3 vs CCR10, we examined the amino-acid composition of their orthosteric binding pockets (**Supp. Fig. 5**). Compared to the more canonical binding pocket composition of CCR3 (many bulky residues conserved among most chemokine receptors), the binding pocket of CCR10 features multiple large-to-small residue substitutions. For example, CCR3 residues W90(2.60), H114(3.33) and Y172(4.65) correspond to Ala98(2.60), Ser121(3.33) Ser180(4.65) in CCR10 (**Supp. Fig. 5**). Additionally, the ECL2 of CCR3 is strongly acidic, featuring several negatively charged residues (e.g., Glu173, Glu175, Glu176, Glu189), whereas the ECL2 of CCR10 is more neutral/basic (e.g., Gln181, Gln184, Arg185, Arg192, Glu197, Gln201) (**Supp. Fig. 5A**, **Fig. 3F right, Fig. 4E right**).

The AlphaFold model of the CCR3-CCL28 complex suggests that both of these factors contribute to the high affinity and potency of CCL28 (but not CCL27) towards CCR3 **(Fig. 4D, E)**. Despite reduced size, compared to CCR10 (**Supp. Fig. 5B, C**), the CCR3 binding pocket favorably accommodates the smaller CCL28 N-terminus in the geometry loosely resembling its complex with CCR10. The side-chain of Ile1 packs underneath the ECL2 of CCR3 and the conserved disulfide bridge Cys106(3.25)-Cys183(ECL2) **(Fig. 4D**). Leu2 of CCL28 does not reach Y113(3.32) in the CCR3 complex (**Fig. 4D**) as it does in the CCR10 complex (**Fig. 3G**). Instead, it is above the residue at position 7.39, which in CCR3 is a Glu (Glu287(7.39)), as in most other chemokine receptors; it is Ser296(7.39) in CCR10. In CRS1, the acidic residues in the proximal N-terminus of CCR3 favorably interact with the basic patches in the CCL28 4_10_ helix and 40s loop (**Fig 4E**, left). A major contribution to the binding energy seems to be the charge-charge interaction between the very basic β_1_-strand and C-terminus of the chemokine on one side, and the acidic ECL2 of the receptor on the other (**Fig 4E**, right).

We also generated a model of CCR3 with NT27-CCL28 (**Fig. 4F**) to illustrate why this chimeric chemokine continues to bind but cannot activate the receptor. The model suggests that the longer N-terminus of CCL27, that also has a bulkier Phe residue in position 1, cannot fit in the smaller binding pocket of CCR3 in the orientation that in other complexes provides activation-associated contacts. Despite this, the CRS1 and ECL2 interactions with this chimera continue to be favorable and likely drive the binding observed experimentally (**Fig. 4C**).

### CCL28 binds GAGs with significantly higher affinity than CCL27, providing another indication of the non-redundant functions of these chemokines

In addition to binding chemokine receptors, chemokines interact with the GAG chains of proteoglycans on cell surfaces and the extracellular matrix as a mechanism to localize them in chemokine gradients that provide directional cues for migrating cells (34, 59, 60). We previously demonstrated that CCL27 interacts weakly with GAGs and has a diffuse GAG binding surface (28), while characterization of CCL28 binding to GAGs has been limited (61). To assess the GAG-binding propensities of CCL28, we employed sepharose heparin (HP) affinity chromatography, which is commonly used to approximate the relative affinities of heparin-binding proteins based on the concentration of NaCl required to elute the proteins from an HP column (62) (**Fig. 5A**). For CCL28, ∼1.2 M NaCl was required for elution, suggesting that it is a potent heparin-binding protein, in stark contrast to CCL27 that showed a much weaker interaction, requiring only ∼680 mM NaCl for elution (**Fig. 5A**). We also evaluated whether the extended C-terminus of CCL28 contributes to GAG binding, similar to CCL21 and CXCL12γ (63, 64). To test this, we generated [1-81]-CCL28, which lacks most of the 28-residue extension, but maintains the unusual third disulfide that links the β1-strand of the globular domain to a region just past the C-terminal α-helix (**Fig. 1A**). The [1-81]-CCL28 truncation mutant eluted at ∼1.1 M NaCl, 143 mM less than that required for WT CCL28 (**Fig. 5A**), indicating that while the C-terminus contributes to GAG binding, the globular domain contributes to the bulk of the GAG-binding affinity consistent with its highly basic surface (**Supp. Fig. 6**). For comparison, we evaluated two other well-characterized chemokines, CCL19, CCL21, and a C-terminal truncation mutant of CCL21, [1-79]-CCL21 (**Fig. 5A**). CCL21 is known to bind GAGs with high affinity and in our hands eluted from the HP column at a NaCl concentration of 956 mM, whereas CCL19 is considered to be a more diffusible, weak-binding chemokine, and eluted at 676 mM NaCl, consistent with prior studies (61, 63, 65, 66). Moreover, removal of the C-terminal tail of CCL21 resulted in a variant with a heparin-binding propensity similar to CCL19, as demonstrated previously (66). Thus, while CCL27 is similar to CCL19 in its GAG binding affinity, CCL28 has an even higher affinity than CCL21; and even after deletion of the CCL28 C-terminal tail, it still retained an affinity equivalent to WT CCL21 ([NaCl]_H_ ∼ 1 M) (**Fig. 5A**).

**Figure 5.**
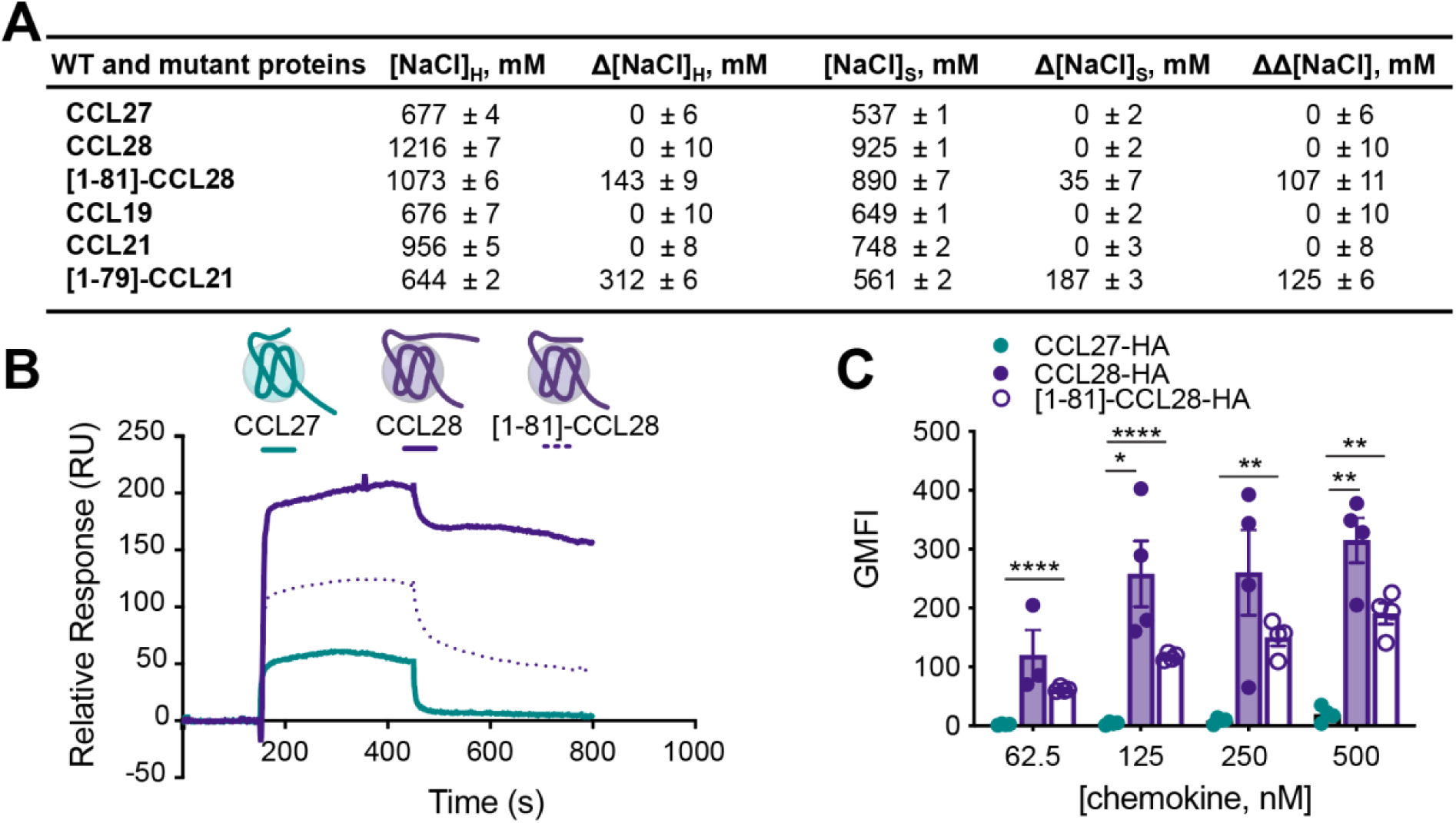
GAG binding characterization of WT and mutant proteins. **(A)** GAG binding characterization of CCL27, CCL28, CCL19, CCL21 and mutants by heparin sepharose chromatography. The concentration of NaCl (mM) required to elute protein from a heparin or a SP sepharose affinity column is reported as [NaCl]_H_ and [NaCl]_S_, respectively. Δ[NaCl] values represent the difference in [NaCl] required to elute mutant versus WT protein (Δ[NaCl]= [NaCl]^WT^ - [NaCl]^mutant^) from either heparin sepharose (Δ[NaCl]_H_) or SP-sepharose (Δ[NaCl]_S_). ΔΔ[NaCl] is a measure of the specificity of interaction, where ΔΔ[NaCl] = Δ[NaCl]_H_ –Δ[NaCl]_S_. A positive specificity index for a mutant means that the difference between the amount of NaCl required to elute the mutant from the heparin-sepharose column as compared to WT was greater than the same difference for the SP-sepharose column, suggesting specificity for heparin beyond electrostatic interactions generally associated with affinity for SP-sepharose. All data represent the mean ± SD of three independent experiments. (**B**) SPR sensorgrams of CCL27, CCL28 and [1-81]-CCL28 showing the resulting signals (relative response in RU) for interaction with immobilized HS with the same concentration of injected chemokine (1000 nM concentration). **(C)** CCL27, CCL28 and [1-81]-CCL28 binding of CHO-K1 cells expressing endogenous HS. C-terminally HA-tagged CCL27, CCL28 or [1-81]-CCL28 was used to determine the extent of chemokine binding to CHO-K1 cells. Cells were incubated with varying concentrations of HA-tagged chemokine for 30 min on ice, washed and the geometric mean fluorescence intensity (GMFI) detected by flow cytometry analysis using an anti-HA antibody normalized to an isotype control. Data shown is the mean ± SEM from single measurements obtained for each concentration combined from independent experiments (n=3-4). Statistical analysis was performed by mixed-effects analysis with *post hoc* Tukey’s multiple comparisons test. (∗∗∗∗p < 0.0001; ∗∗p < 0.01; ∗p < 0.05; CCL27-HA : CCL28-HA at 62.5 nM, p = 0.1915; CCL28-HA : [1-81]-CCL28-HA at 62.5 nM, p = 0.4916; CCL28-HA : [1-81]-CCL28-HA at 125 nM, p = 0.1659; CCL27-HA : CCL28-HA at 250 nM, p = 0.0798; CCL28-HA : [1-81]-CCL28-HA at 250 nM, p = 0.4024; CCL28-HA : [1-81]-CCL28-HA at 500 nM, p = 0.0812).

Affinity values and association/dissociation rates from Surface Plasmon Resonance (SPR) experiments supported the above conclusions (**Fig. 5B, Supp. Fig. 7**). For the SPR experiments, biotinylated heparan sulfate (HS) was immobilized on neutravidin-coated BIAcore C1 chips and varying concentrations of WT or mutant chemokine flowed over the chip, as previously described (67, 68) (**Supp. Fig. 8**). In qualitative agreement with the HP affinity chromatography data, the SPR results revealed an ∼1000-fold difference in the affinity of CCL27 and CCL28 for HS (3.2 μM vs. 3 nM, respectively, (**Fig. 5B, Supp. Fig. 7**). The C-terminal truncation mutant, [1-81]-CCL28, exhibited only slightly weaker affinity for HS relative to WT CCL28 (15 nM vs. 3 nM) in contrast to the complete loss-of-binding observed for the CCL21 C-terminal truncation mutant, [1-79]-CCL21 (**Fig. 5B, Supp. Fig. 7**).

To determine whether the heparin-binding propensities of CCL27 and CCL28 correlate with their ability to bind cell surface GAGs, we incubated C-terminally HA-tagged CCL27, CCL28 or [1-81]- CCL28 with CHO-K1 cells, which endogenously express HS, and measured bound chemokine by flow cytometry using an anti-HA antibody (**Fig. 5C**). These data revealed substantial accumulation of CCL28 and diminished but still appreciable interaction of [1-81]-CCL28, consistent with the HP sepharose and SPR data (**Fig. 5A, B, Supp. Fig. 7**). Together, these data highlight additional non-redundant properties of two chemokines that share CCR10 as their receptor.

## Discussion

CCL27 and CCL28 are both involved in epithelial immunity through their common receptor CCR10, but in different capacities: CCL27 is primarily associated with skin where it attracts T cells, while CCL28 is expressed by mucosal tissue and recruits antibody-producing B lymphocytes (8, 69, 70). Given their clear functional differences, prior studies have sought to determine if the ligands differ in their signaling bias, i.e., their ability to preferentially activate G protein vs β-arrestin pathways downstream of CCR10 (71, 72). However, there has been little in the way of structure-function studies, which can provide insight into activation mechanisms and lead to the discovery of variants with altered pharmacological properties.

In this study, we investigated how CCL27 and CCL28 activate CCR10 with a combination of N-terminal mutations, deletions and chimeric swaps of the chemokines, coupled with AlphaFold modeling of chemokine complexes with CCR10. Key findings include the discovery that the first two amino acids (Phe1 and Leu2) of CCL27 are critical for its ability to activate CCR10. AlphaFold models suggest that these two residues trigger CCR10 activation upon interaction with residues at positions 3.32 and 7.39 in the receptor binding pocket that are known to be involved in the activation of homologous receptors receptors (49–53). Notably, the two-residue deletion mutant resulted in an antagonist variant ([3-88]-CCL27) that may provide the starting point for the rational design of higher affinity antagonists that could have therapeutic value. For example, CCL27 has previously been implicated in atopic dermatitis due to its ability to recruit CCR10 expressing CD4+ and CD8+ T cells; moreover, administration of an anti-CCL27 neutralizing antibody reduced dermal inflammation along with T and Mast cell accumulation, T cell proliferation and the presence of inflammatory cytokines (15). CCR10 has also been implicated in rheumatoid arthritis although the efficacy of inhibiting the receptor on disease pathogenesis remains to be determined (73). Numerous reports have described a role of CCR10 and its ligands in cancer although the benefits of CCR10 antagonism may be cancer dependent: the receptor can have pro-tumoral effects when expressed on regulatory T cells or tumor cells or anti-tumoral effects when expressed on CD4+ and CD8+ T cells (74–77). Thus, [3-88]-CCL27, or higher affinity analogues, may be useful for further probing the efficacy of inhibiting CCR10 for the treatment of these diseases and potentially used directly as protein therapeutics.

We also showed that N-terminal extension of CCL27 with a Phe results in a superagonist that is substantially more potent (10-fold) and efficacious (1.1-fold) than WT CCL27 in promoting cell migration in a bare filter assay, with even more enhanced effects observed in inducing transendothelial cell migration and β-arrestin recruitment. An AlphaFold model of the CCR10:F-CCL27 complex suggests this is due to the extension of Phe0 into the major subpocket of the receptor and direct interactions with a hydrophobic stack of residues in activation helix 6 in a manner that influences the "toggle switch" residue at position 6.48. The Phe addition may also result in greater stabilization of the active receptor conformation and/or increased residence time of the ligand on the receptor; however, the CCR10:CCL27 system proved recalcitrant to traditional radioligand binding assays, and therefore affinities and off-rates were not determined. Regardless of mechanism, F-CCL27 may be useful as an adjuvant to enhance the efficacy of vaccines: for example, plasmids encoding CCL27 along with the SARS-CoV-2 spike protein resulted in increased anti-spike protein antibodies to CD8+ T cells at mucosal surfaces, which conferred significantly enhanced immunity against SARS-CoV-2 (78). Both CCL27 and CCL28 have also been shown to provide immunity against influenza and HIV (30, 31, 79) due to the fact that adjuvant activity is dependent on CCR10-expressing plasma B cells that produce IgA and accumulate at mucosal surfaces (14). However, whether superagonism conferred by F-CCL27 results in a more potent response *in vivo* remains to be determined.

In contrast to CCL27, extension of the N-terminus of CCL28 with a Phe (or Phe-Phe) did not enhance its ability to promote CCR10-mediated cell migration. Nevertheless, the impact of swapping the N-termini of CCL28 and CCL27 on CCR10-mediated migration and internalization suggests that the CCL28 N-terminus is a stronger driver of agonist promoted responses. And in fact, the chimera (NT28-CCL27) also proved to be a superagonist of CCR10 that like F-CCL27 may have therapeutic utility. In addition to having potential as more potent vaccine adjuvants than the WT ligands, superagonists may also serve as "functional antagonists" if they enhance receptor internalization, as we observe for F-CCL27 and NT28-CCL27, especially if internalization results in trafficking to degradation pathways and/or inhibition of receptor recycling (80–82). Prior examples of chemokine-based functional antagonists include AOP-RANTES/CCL5 and PSC-RANTES/CCL5, both of which inhibit HIV entry into cells by promoting internalization and intracellular retention of CCR5 (56, 83).

In addition to identifying variants of CCL27 with distinct pharmacological properties, AlphaFold models of complexes of CCR10 and CCR3 with CCL27 and CCL28 also provided insight into the ability of CCR10 to be activated by both CCL27 and CCL28 and the specificity of CCR3 for CCL28. Specifically, the orthosteric pocket of CCR3 is too small for the longer/bulkier N-terminus of CCL27 but accommodates that of CCL28. CCL28 (pI=10.35) is also more basic than CCL27 (pI=9.1) (**Supp. Fig. 6**) and shows electrostatic complementarity with the acidic N-terminus and ECL2 of CCR3, which is lacking in CCL27. The more basic nature of CCL28 translates into its higher affinity for heparin, HS and cell surface GAGs compared to CCL27, which may be critical for the accumulation of CCL28 in mucosal tissues in the context of its chemoattractant functions. For example, HS within the mouse caecum (colon) is expressed at low levels in naïve tissue and upregulated at the base of the crypts following chronic infection (84), which may benefit from the intrinsic high-affinity (and slow off-rate) HS-binding properties of CCL28 for directed immune cell migration to focused regions of HS expression. On the other hand, expression of HS GAG on the skin is localized to the vasculature, immune cell surface and epidermis at rest and following inflammation (85), perhaps providing sufficient sites of retention for the more weakly HS-binding CCL27. Similar to defensins and histatin (86, 87), the abundance of positively charged residues is also important for the broad-spectrum antimicrobial activity of CCL28, which facilitates disruption of bacterial membranes (86). Thus, CCL28 has been described as having a dual role in mucosal protection through its ability to recruit CCR3- and CCR10-expressing cells and its antimicrobial activity (87).

Altogether, the data in this study provide mechanistic explanations for the non-redundant biological functions of CCL27 and CCL28 along with structural insights that account for their receptor selectivity and distinct context-dependent signaling. The pharmacologically unique antagonist and superagonist variants discovered in the course of our studies may prove useful for interrogating the value of targeting CCR10 for specific diseases.

## Experimental procedures

### Chemokine expression and purification

WT and mutant CCL27 and CCL28 and CCL7-HA were expressed as a His-ubiquitin fusion using the pHUE vector and purified from inclusion bodies according to established procedures (28, 67). CCL8 was expressed and purified with a N-terminal His(8x)-tag followed by an enterokinase cleavage site, similar to previously established procedures (88). Protein identity and purity was confirmed by electrospray ionization mass spectrometry (Scripps Center for Metabolomics or the UC San Diego Molecular Mass Spectrometry Facility). Specified mutants were generated by QuikChange site-directed mutagenesis (Stratagene) and purified in the same manner as WT protein.

### Cell culture

Murine L1.2 pre-B cells stably expressing human CCR10 (kind gift of Eugene Baker, Stanford University, USA) or human CCR3 (generated in-house through limiting dilution) were maintained in RPMI 1640 media (Gibco) supplemented with 10% fetal bovine serum (FBS) (Gibco), 1% MEM non-essential amino acids, 1% sodium pyruvate, 0.1% BME, and 300 µg/mL geneticin and grown at 37°C with 5% CO_2_. EA926 human umbilical vein endothelial cells (kind gift of the Shyy lab, UCSD, USA) were cultured in DMEM/10% FBS and grown at 37°C with 5% CO_2_. HEK293 cells (ATCC, cat# CRL-1573) were maintained in DMEM/10% FBS and grown at 37°C with 5% CO2. Mycoplamsa testing was performed using either the MycoStrip^TM^ (InvivoGen) or MycoAlert Plus (VWR) mycoplasma detection kit, with no mycoplasma detected.

### DNA plasmids and cloning

CCR10-RLucII was generated by subcloning the CCR10 coding sequence into a vector containing the RLucII tag. The βarr2-GFP10 construct was kindly gifted by Nikolaus Heveker (Université de Montréal, Canada).

### Chemokine binding on CHO cells

Binding of C-terminally HA-tagged chemokine on CHO-K1 cells (kind gift of Dr. Jeffrey Esko, UC San Diego, CA, USA) was performed essentially as described previously (67) with a few minor modifications. Cells were treated with HA-tagged WT or mutant chemokine (62.5-500 nM) prepared in FACs buffer (PBS + 0.5% BSA) and incubated on ice for 30 min. Unbound chemokine was removed by washing cells three times with FACs buffer. Chemokine levels were then detected by incubation with a PE-conjugated anti-HA monoclonal mouse antibody (Miltenyi Biotec, cat# 130-092-257) or PE-conjugated mouse isotype control (BD, cat #555787), according to manufacturer instructions and analyzed by a Guava EasyCyte 8HT flow cytometer (EMD Millipore). Post-acquisition analysis with FlowJo (Tree Star, Inc) was used to determine the geometric mean fluorescent intensity for each sample normalized to isotype control; data are plotted as the mean ± SD of at least three independent experiments.

### Bare filter and transendothelial cell migration assays

Traditional bare filter migration assays were performed as described previously (67, 68) with some modifications. Briefly, cells were resuspended at 1.5 - 2.5 x 10^6^ cells/mL in RPMI 1640 + 10% FBS and 100 µL of cells were added to the upper chamber of a 5 µm pore size 24-well transwell filter insert (Corning), or 75 µL of cells added to the upper chamber of a 96-well transwell setup (Corning). Cells were allowed to migrate towards varying concentrations of CCL27 or CCL28 present in the lower chamber. After 2 h of incubation at 37°C/5% CO_2_, migrated cells were counted on a Guava EasyCyte 8HT flow cytometer (EMD Millipore) by counting the number of cells present in 30 s. Migration was calculated as the percent of migration compared to the maximal number of possible cells migrated (no filter). To determine whether chemokine variants retained their ability to bind to surface receptor, competition binding migration assays were also employed. In these instances, bare filter migration assays were performed similar to above; however, each well contained 50 nM WT CCL27 in addition to increasing concentrations of mutant CCL27. While these assays do not allow the quantification of binding constants, they can indicate whether a chemokine mutant retains the ability to interact with receptor in a manner competitive with WT chemokine. Lastly, transendothelial migration assays were performed identically to the bare filter assays, with the exception that filters were first coated with 0.1 mg/mL type I collagen (Purecol, Advanced Biomatrix) for 1 h at 37°C and then EA926 cells were added to each well (0.1 x 10^6^ cells/filter in 100 µL) and allowed to grow to confluency for 2 days at 37°C with 5% CO_2_. After 2 days, transendothelial migration assays were performed as described for the bare filter experiments. Assays were undertaken in duplicate or triplicate with at least three independent experiments performed for each dataset. Representative data are plotted as the mean ± SD, unless otherwise stated.

### Chemokine competition binding assays

Competition binding experiments were performed on L1.2/CCR3 cells using C-terminally HA-tagged CCL7 (CCL7-HA). CCR3 expression was induced by incubation of cells with 5 mM sodium butyrate for ∼22h prior to conducting the experiment. Cells were then incubated with 250 nM of CCL7-HA alone or in combination with a 4-fold excess of competing ligand (1000 nM) for 30 min on ice, cells were washed and then CCL7-HA levels were determined by flow cytometry analysis using an anti-HA antibody. Results shown are the mean ± SD of single measurements obtained for each concentration combined from three independent experiments; the dotted line depicts 100% retention of bound CCL7-HA.

### Heparin-sepharose binding assays

The amount of NaCl required to elute CCL27, CCL28, CCL19, CCL21 and mutants from a 1 -mL HiTrap Heparin HP column (GE Healthcare) or a 1-mL HiTrap SP HP column (GE Healthcare) was determined as previously described (68). Each assay was performed in triplicate; data is plotted as the mean ± SD.

### Surface Plasmon Resonance

A BIAcore 3000 instrument (GE Healthcare) was used to perform SPR using a C1 SPR chip surface coated with HS, as described previously (67, 68). Briefly, paired cells on a C1 SPR chip (GE Healthcare) were equilibrated in running buffer (10 mM HEPES, 150 mM NaCl, 3 mM EDTA, 0.05% Tween-20, pH 7.4) before activation with NHS:EDC (1:1 mixture) and immobilization of neutravidin (0.2 mg/ml), until saturation, in 20 mM Sodium Acetate, pH 6.0, excess neutravidin was removed by washing with regeneration buffer (0.1 M Glycine, 1 M NaCl, 0.1% Tween-20, pH 9.5). Remaining active sites were blocked by flowing 1 M ethanolamine over the paired cells. Biotinylated HS (0.2 mg/ml) (67) was then flowed over one of the paired flow cells followed by extensive washing with running and regeneration buffer. Chemokines, at the indicated concentrations, were then passed over these paired cells in running buffer (40 μL/min, to limit mass transfer effects) and the HS specific response monitored by subtracting the signal from the paired flow cell without HS from that with HS. The surface was regenerated between chemokine injections using regeneration buffer. The resulting sensorgrams were then analyzed using BIAevaluation software and apparent dissociation constants (*K*_D_) were obtained using a 1:1 Langmuir interaction model and/or steady-state analyses where possible (67, 89). Fitting of the data was assessed visually and using the Chi^2^ values, where a Chi^2^<10 was considered to be indicative of a good fit. In instances where HS-bound chemokine signal reached saturation during injection, at a sufficient range of concentrations, these maximum signal values were plotted against concentration and analyzed with the 1:1 steady-state affinity model equilibrium analysis in the BIAevaluation software (GE Healthcare). The resulting affinities are considered to be “apparent affinities” due to the issues associated with analyzing chemokine:glycosaminoglycan interactions discussed in detail elsewhere (62, 67).

### Calcium flux assays

Chemokine-mediated calcium flux was measured using the Screen Quest^TM^ Fura-2 No Wash Calcium Assay Kit (AAT Bioquest). Briefly, 100 µL of 2 x 10^5^ L1.2/CCR10 cells suspended in HHBS (1X Hank’s Balanced Salt Solution with 20 mM HEPES, pH 7.2) were seeded in a black-wall, clear-bottom assay plate. The plate was centrifuged at 140 x g for 5 min at 25°C and incubated for 30 minutes at 37°C with 5% CO_2_. Then 100 µL of Fura-2 AM dye-loading solution prepared in HHBS was added to the cells, followed by one-hour incubation at 37°C with 5% CO_2_. After that, the plate was loaded to the TECAN Spark multimode microplate reader for receptor desensitization measurements. Chemokine samples (10 µL in HHBS with 1% bovine serum albumin) were injected to the wells of interest by the Te-Inject™ module after 10th and 75th reading cycles. All assays were performed at 37°C and monitored by fluorescence intensity (excitation: 340/380 nm; emission: 510 nm). Relative Fluorescence Units (RFUs) were calculated by the ratio of 510-nm emission in 340- and 380-nm excitation and normalized to baselines values from the first 10 cycles. The processed data was plotted using GraphPad Prism (GraphPad Software).

### Receptor internalization assays

Internalization assays were performed as described previously (90), with some modifications. Briefly, L1.2 cells expressing CCR10 were prepared at 1 x 10^6^ cells/mL in assay buffer (RPMI + 10% FBS) and chilled on ice in a 96-well plate. Cells were spun down to remove assay buffer and resuspended in 100 µL (100,000 cells/sample) of varying concentrations (3.1 to 100 nM) of WT or mutant chemokine prepared in cold assay buffer followed by incubation at 37°C/5% CO_2_ for 45 min. Cells were then diluted in cold FACs buffer and washed twice with FACs buffer. CCR10 surface levels were measured by staining cells with an anti-hCCR10 PE-conjugated rat monoclonal antibody (R&D, cat# FAB3478P) or isotype control according to manufacturer instructions. Data was acquired on a Guava EasyCyte 8HT flow cytometer (EMD Millipore) and analyzed with FlowJo software (Treestar, Inc.). Data was plotted using GraphPad Prism (GraphPad Software) and shows the percent of CCR10 remaining on the cell surface (compared to the maximal CCR10 levels detected with no chemokine added) from three independent experiments (mean ± SD).

### BRET assays

For evaluation of β-arrestin recruitment to CCR10, HEK293 cells were transfected with Mirus TransIT-Lt1 reagent when cells reached ∼70% confluency. Transfection included 100 ng of BRET donor (CCR10-RlucII) and 2 μg of BRET acceptor (βarr2-GFP10). Assays were performed ∼48 h post-transfection. On the day of the assay, cells were seeded into 96-well plates (100,000 cells per well) and incubated in Tyrode’s Buffer (140 mM NaCl, 12 mM NaHCO_3_, 5.6 mM D-Glucose, 2.7 mM KCl, 1 mM CaCl_2_, 0.5 mM MgCl_2_, 0.37 NaH_2_PO_4_, 25 mM HEPES) for 40 min followed by the addition of the substrate, Prolume Purple (NanoLight Technologies) to a final concentration of 5 μM. After 10 min, baseline BRET measurements were performed to measure donor and acceptor emissions at their respective wavelengths: 480 nm and 530 nm. Subsequently, the indicated concentrations of chemokines were added to wells and BRET measurements were taken every 5 min for 30 min. The average time of each reading was calculated, and raw BRET values were converted into a BRET ratio (donor/acceptor). Data was plotted using GraphPad Prism (GraphPad Software) and represented as the inverse of the BRET ratio and normalized to the maximal WT response. All data points are the mean ± SD from a minimum of three independent experiments performed in at least duplicates.

### Molecular modeling

Structural models of CCR3 and CCR10 complexes with WT and mutant chemokines were constructed by AlphaFold2 Multimer v2.3.2 (91, 92) locally installed on the UCSD Triton Shared Computing Cluster (TSCC). The following amino-acid sequences were used for modeling:

~~~
CCR10:
MGTEATEQVSWGHYSGDEEDAYSAEPLPELCYKADVQAFSRAFQPSVSLTVAALGLAGNGLVLATHLAARRAARSPTSAHLLQLALADLLLALTLPFAAAGALQGWSLGSATCRTISGLYSASFHAGFLFLACISADRYVAIARALPAGPRPSTPGRAHLVSVIVWLLSLLLALPALLFSQDGQREGQRRCRLIFPEGLTQTVKGASAVAQVALGFALPLGVMVACYALLGRTLLAARGPERRRALRVVVALVAAFVVLQLPYSLALLLDTADLLAARERSCPASKRKDVALLVTSGLALARCGLNPVLYAFLGLRFRQDLRRLLRGGSCPSGPQPRRGCPRRPRLSSCSAPTETHSLSWDN
~~~

~~~
CCR3:
MTTSLDTVETFGTTSYYDDVGLLCEKADTRALMAQFVPPLYSLVFTVGLLGNVVVVMILIKYRRLRIMTN IYLLNLAISDLLFLVTLPFWIHYVRGHNWVFGHGMCKLLSGFYHTGLYSEIFFIILLTIDRYLAIVHAVF ALRARTVTFGVITSIVTWGLAVLAALPEFIFYETEELFEETLCSALYPEDTVYSWRHFHTLRMTIFCLVL PLLVMAICYTGIIKTLLRCPSKKKYKAIRLIFVIMAVFFIFWTPYNVAILLSSYQSILFGNDCERSKHLD LVMLVTEVIAYSHCCMNPVIYAFVGERFRKYLRHFFHRHLLMHLGRYIPFLPSEKLERTSSVSPSTAEPE LSIVF
~~~

~~~
CCL27
FLLPPSTACCTQLYRKPLSDKLLRKVIQVELQEADGDCHLQAFVLHLAQRSICIHPQNPSLSQWFEHQERKLHGTLPKLNFGMLRKMG
~~~

~~~
CCL28
ILPIASSCCTEVSHHISRRLLERVNMCRIQRADGDCDLAAVILHVKRRRICVSPHNHTVKQWMKVQAAKKNGKGNVCHRKKHHGKRNSNRAHQGKHETYGHKTPY
~~~

~~~
F-CCL27:
FFLLPPSTACCTQLYRKPLSDKLLRKVIQVELQEADGDCHLQAFVLHLAQRSICIHPQNPSLSQWFEHQERKLHGTLPKLNFGMLRKMG
~~~

~~~
NT28-CCL27:
ILPIASSCCTQLYRKPLSDKLLRKVIQVELQEADGDCHLQAFVLHLAQRSICIHPQNPSLSQWFEHQERKLHGTLPKLNFGMLRKMG
~~~

~~~
NT27-CCL28:
FLLPPSTACCTEVSHHISRRLLERVNMCRIQRADGDCDLAAVILHVKRRRICVSPHNHTVKQWMKVQAAKKNGKGNVCHRKKHHGKRNSNRAHQGKHETYGHKTPY
~~~

An ensemble of 25 models (5 random seeds with 5 models per seed) was built for each of the following complexes:

- CCR10 w/ CCL27
- CCR10 w/ F-CCL27
- CCR10 w/ NT28-CCL27
- CCR10 w/ CCL28
- CCR10 w/ NT27-CCL28
- CCR3 w/ CCL28
- CCR3 w/ NT27-CCL28

Using ICM software version 3.9-2e (93) the chemokines were modified to include a free positively charged N-terminus (NH_3_^+^). For each of the models, receptor interactions with the chemokine N-terminus and 30s loop were additionally refined in ICM in two stages: one employing 3D grid potentials and another full-atom representation of all components. During the first stage, the receptor binding pocket was represented with a set of grid interaction potentials, including those for van der Waals, electrostatic, hydrogen bonding and apolar surface interactions (94, 95). The N-terminus and the 30s loop of the chemokine were built *ab initio* with the following sequences:

- **CCL27**: Phe1-LLPPSTAC-Cys10 and Glu33-ADGDCHL-Gln41
- **CCL28**: Ile1-LPIASSC-Cys9 and Arg31-ADGDCDL-Ala39
- **F-CCL27**: Phe0-FLLPPSTAC-Cys10 and Glu33-ADGDCHL-Gln41
- **NT28-CCL27**: Ile1-LPIASSC-Cys9 and Glu32-ADGDCHL-Gln40
- **NT27-CCL28**: Phe1-LLPPSTAC-Cys10 and Arg32-ADGDCDL-Ala40

An explicit disulfide bond was imposed between the first cysteine of the chemokine N-terminus and the only cysteine in the 30s loop; terminal residues of each chemokine fragment were tethered to the corresponding residues in the template. The conformational stack of each system was populated from the AlphaFold2 model ensemble, and the system was then thoroughly sampled in the receptor potential grids, using biased probability Monte Carlo sampling in ICM, to optimize and expand on this conformational stack. For the second stage, the resulting conformational stack was merged with full atom models of the receptor, and at least 10^8^ steps of Monte Carlo optimization were performed, allowing for the same level of flexibility in the chemokine fragments with added full flexibility of receptor binding pocket sidechains. For full-atom sampling, van der Waals, torsional, hydrogen bonding, electrostatic, and disulfide bond energy terms were used. The resulting conformations were ordered by predicted energy; top ranking conformations were used for visualization and analysis of molecular interactions.

### Binding free energy calculations

Prime 6.9 module of Schrödinger suite was used to perform molecular mechanics generalized Born and surface area solvation (MM-GBSA) ((54, 55), Schrödinger, LLC, New York, NY, 2021). Binding free energy (**Δ**G) for WT or mutant chemokines in complex with receptor was calculated as follows:

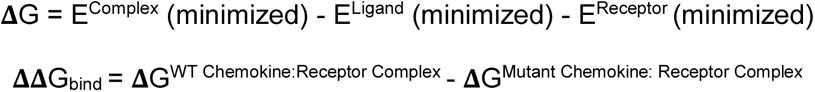

Systems were performed with the VSGB 2.1 model for implicit solvation (96) and OPLS4 force field with water as solvent. Residues within 5 Å of the chemokine atoms were treated as flexible and subject to minimization. Relative binding energies are reported in kcal/mol.

## Data availability

All data needed to evaluate the conclusions in the study are present in the manuscript, Supplementary Materials, and Supplementary Data.

## Supporting information

This article contains supporting information.

## Acknowledgements

We thank David Aguilar for cloning the NT28-CCL27, Nicholas Chimileski and Brian Nguyen for cloning the L-CCL27 plasmid and Tetsuya Kawamura for the preparation of some of the chemokines used in these studies.

## Author contributions

Conceptualization: CLS, TMH

Data Curation: MH, AFC, RC, DPD, ALJ, IK, CLS, TMH

Formal Analysis: MH, AFC, RC, DPD, ALJ, IK, CLS, TMH

Funding Acquisition: IK, TMH

Investigation: MH, AFC, RC, DPD, ALJ, CLS

Methodology: MH, AFC, RC, DPD, ALJ, IK, CLS, TMH

Project Administration: IK, CLS, TMH

Resources: MH, AFC, RC, DPD, ALJ, IK, CLS, TMH

Software: RC, IK

Validation: MH, AFC, DPD, ALJ, CLS

Visualization: MH, AFC, RC, DPD, IK, CLS,

TMH Writing – original draft: CLS, IK, TMH

Writing – review and editing: MH, AFC, RC, DPD, ALJ, IK, CLS, TMH

## Funding and additional information

This work was supported by National Institute of Health grants R01 AI37113 and R21 AI121918 to TMH, R01 GM136202, R01 AI161880 and R01 AI118985 to TMH and IK, R21 AI149369 and R21 AI156662 to IK.

## Abbreviations

AF2: AlphaFold 2
BRET: bioluminescence resonance energy transfer
CRS: chemokine recognition site
ECL: extracellular loop
FBS: fetal bovine serum
HS: heparan sulfate
MM-GBSA: molecular mechanics generalized Born and surface area solvation
SPR: surface plasmon resonance
TM: transmembrane

