## Supplemental figures for "Rules of engagement: determinants of chemokine receptor activation and selectivity by CCL27 and CCL28"

**Supplemental figure legends**

**Supplemental Figure 1. Calcium flux analysis of CCL27 and F-CCL27.** (**A-C**) Chemokine-mediated desensitization of L1.2/CCR10 cells as measured by calcium flux. Cells were treated with an initial dose of 20 nM WT or F-CCL27 (injection 1) followed by a second addition of 20 nM F-CCL27 and WT CCL27 (injection 2), respectively, after the 75th reading cycle (at approximately 320 s). Shown are results from three independent experiments performed in technical triplicates.

**Supplemental Figure 2. Cell migration analysis of CCL28 N-terminal mutants.** Cell migration analysis with L1.2/CCR10 cells across varying chemokine concentrations of CCL28, F-CCL28, FF-CCL28 and F-NT27-CCL28. Data are plotted as the percent of cells migrated after a 2 h incubation at 37°C. Results shown are representative data (mean ± SD) from three independent experiments performed in technical triplicates.

**Supplemental Figure 3. β-arrestin recruitment by BRET.** Cells were transiently transfected with CCR10-RlucII and βarr2-GFP10 and then stimulated with varying concentrations of CCL27 or NT28-CCL27. The BRET ratios are shown as a % of WT-CCL27. Results shown are the mean ± SD of at least three independent experiments performed in technical duplicates or triplicates.

**Supplemental Figure 4. Comparison of the conformations and secondary structures of CCL27 (teal) and CCL28 (purple) N-termini in the binding pocket of CCR10.** (**A**) Side view (parallel to the plane of the membrane), the optimal ligand atom placement surface in the receptor's orthosteric binding pocket is shown as a grey mesh. (**B**) Top view (perpendicular to the membrane from the extracellular side), the receptor is shown as a white ribbon.

**Supplemental Figure 5. Comparison of CCR3 and CCR10 binding pockets.** (**A**) Sequence alignment of pocket residues of CCR3 and CCR10. (**B-C**) Top view of CCR10 (**B**) and CCR3 (**C**) orthosteric binding pockets. The optimal ligand placement surface is shown as a mesh. The amino-acid residues that are not conserved between the two receptors are shown as spheres.

**Supplemental Figure 6. Comparison of electrostatic properties of CCL27 (A) and CCL28 (B).** Chemokines are shown in two orientations ('front' and 'back') as surface meshes colored by electrostatic potential).

**Supplemental Figure 7. Binding analysis of chemokines and mutants with HS.** (**A**) Summary of SPR analysis including the association ($k_a$), dissociation ($k_d$) and resulting affinities ($K_D=k_d/k_a$), and steady state affinity analysis where possible, for chemokine and mutant interactions with HS immobilized on the surface of a C1 chip. Chi$^2$ values are included as a measurement of the quality of the data, as described in Experimental Procedures. (**B**) SPR sensorgrams of CCL19, CCL21 and [1-79]-CCL21 showing the resulting signals (relative response in RU) for interaction with immobilized HS with the same concentration of injected chemokine (200 nM).

**Supplemental Figure 8. SPR interaction analysis of chemokines with HS**. (**A**) CCL27 (1000, 750, 500 and 400 nM), (**B**) CCL28 (1000, 750, 500, 400, 250 (repeated), 200, 100 and 50 nM), (**C**) [1-81]-CCL28 (1000, 750, 500, 400, 250 (repeated), 200, 100 and 50 nM), (**D**) CCL19 (200, 150, 100, 75 and 50 nM), (**E**) CCL21 (200, 150, 100, 75, 50, 37.5 and 25 nM) and (**F**) [1-79]-CCL21 (200, 150, 100, 75, 50, 37.5 and 25 nM) with HS immobilized on a C1 chip. Varying concentrations of chemokines were passed over the HS surface and resulting interactions were analyzed using the 1:1 Langmuir model to generate binding affinities (**Supp. Fig. 8**). Shown are the experimental (black lines) and fitted data (pink lines) against the relative response signal

(RUs) for each chemokine concentration series. Where applicable, steady state analysis was also performed (**Supp. Fig. 5A**). (**B, E**) The insets show a plot of the RUs of chemokines at varying concentrations, which was used to determine the binding affinity by steady state analysis.

**Supplemental Figure 1.**

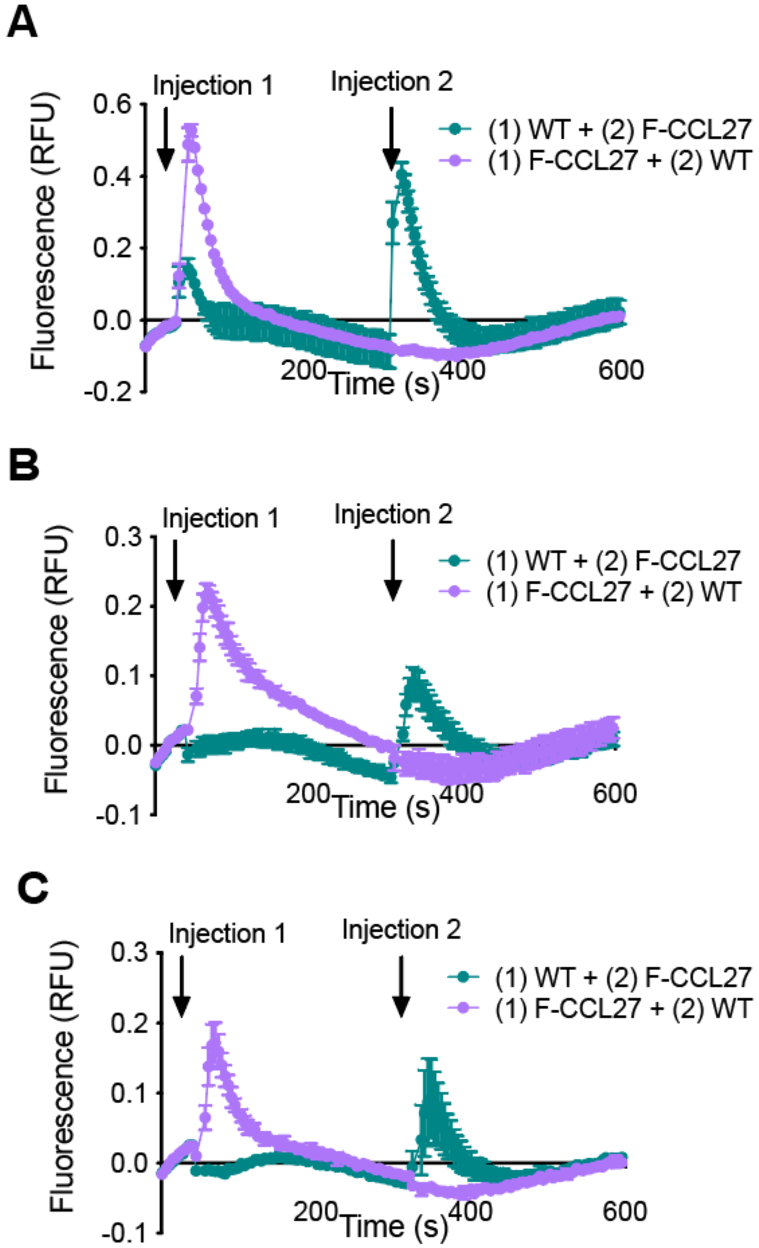

**Supplemental Figure 2.**

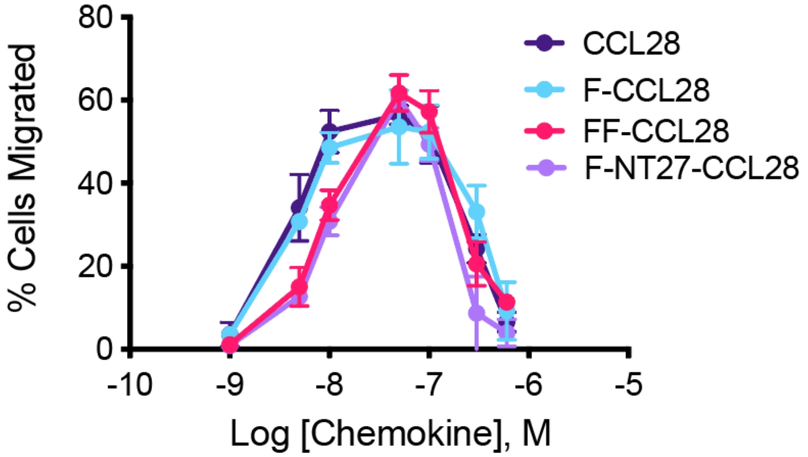

**Supplemental Figure 3.**

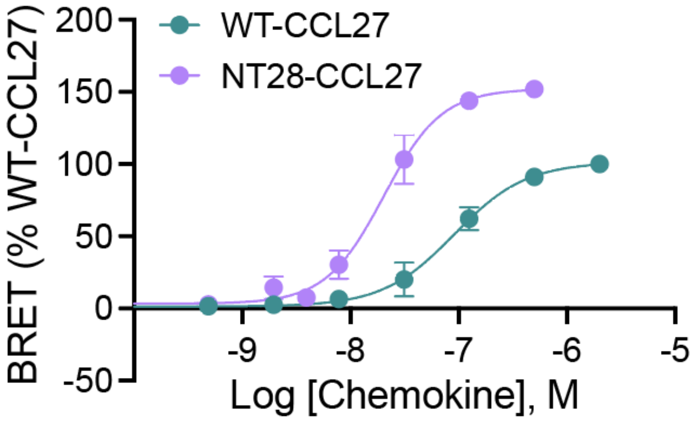

**Supplemental Figure 4.**

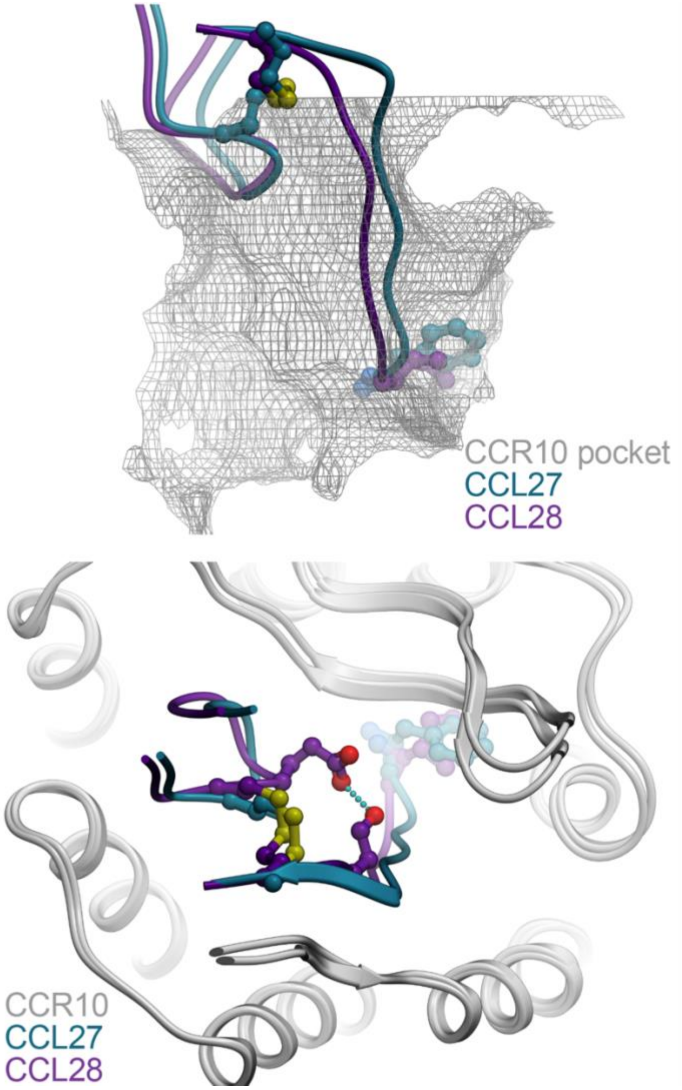

**Supplemental Figure 5.**

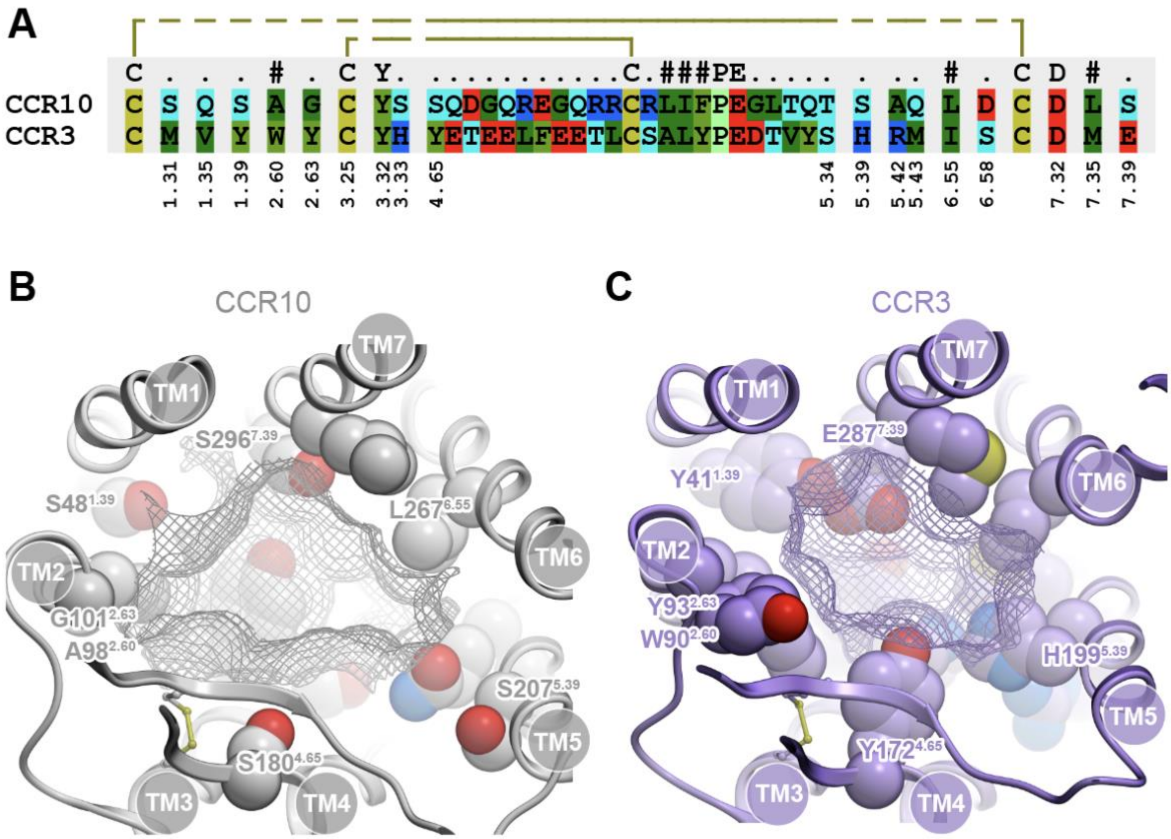

**Supplemental Figure 6.**

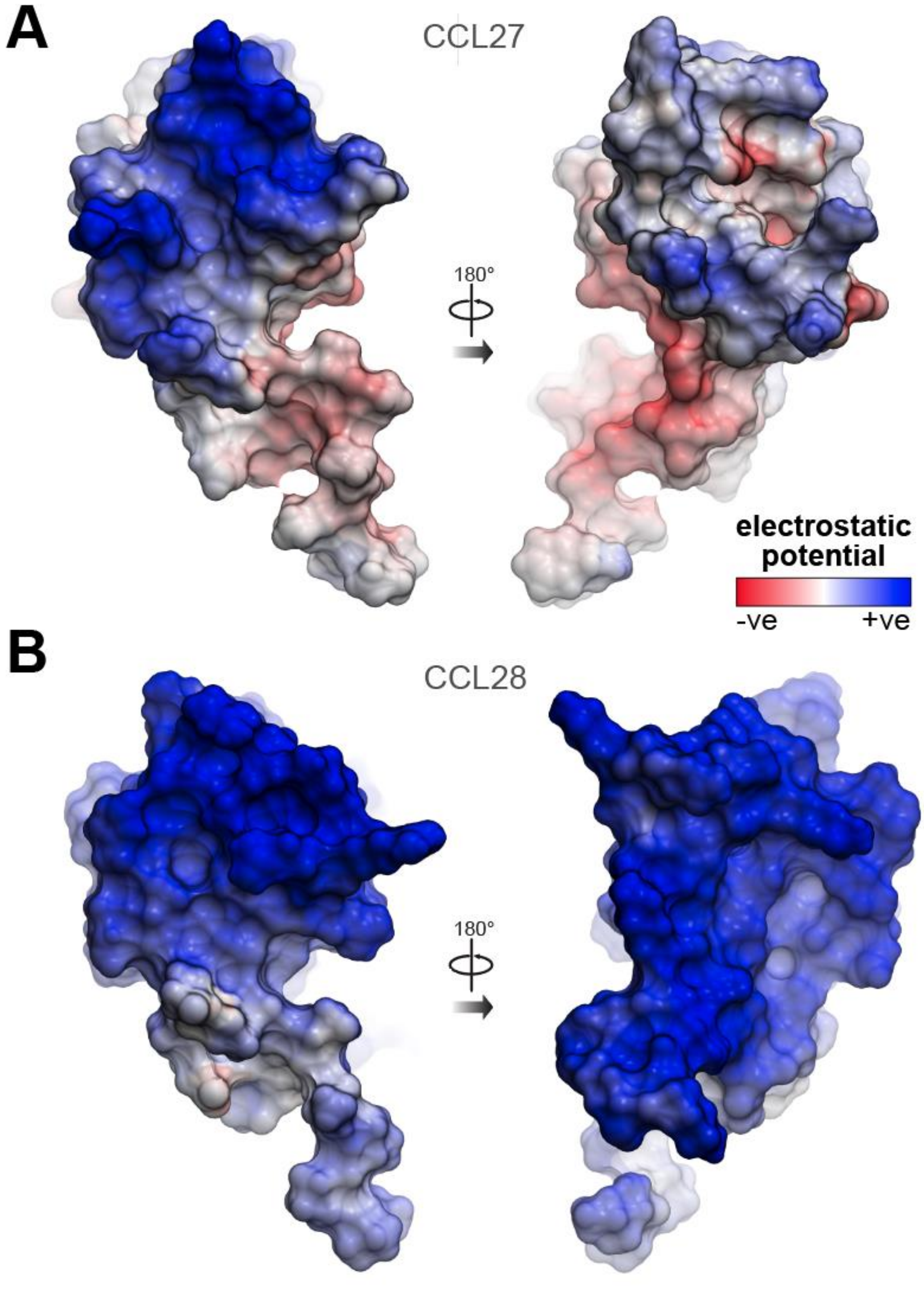

**Supplemental Figure 7.**

**A**

| Chemokine | $k_a$ ($M^{-1}$ $s^{-1}$) | $k_d$ ($s^{-1}$) | $K_D$ (nM) | $Chi^2$ | Steady State Affinity (nM) |
|---|---|---|---|---|---|
| | Heparan Sulfate | | | | |
| **CCL19** | $4.4 \times 10^4$ | $5.0 \times 10^{-3}$ | 113.6 | 3.5 | NPA[a] |
| **CCL21** | $4.1 \times 10^5$ | $7.7 \times 10^{-3}$ | 18.8 | 16.5 | 71 |
| **[1-79]-CCL21** | NOI[b] | NOI | NOI | NOI | NOI |
| **CCL27** | $4.3 \times 10^5$ | $1.4 \times 10^{-1}$ | 3255 | 7.8 | NPA |
| **CCL28** | $3.1 \times 10^5$ | $9.4 \times 10^{-4}$ | 3 | 78.2 | 71 |
| **[1-81]-CCL28** | $1.7 \times 10^5$ | $2.6 \times 10^{-3}$ | 15.3 | 21.1 | 228 |

[a]NPA- no possible analysis

[b]NOI- no observable interactoin

**B**

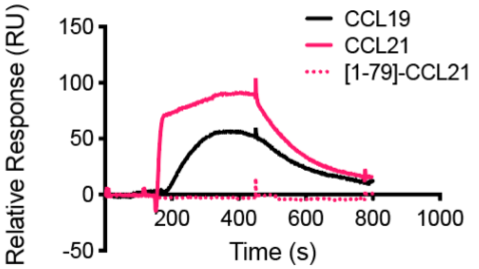

**Supplemental Figure 8.**

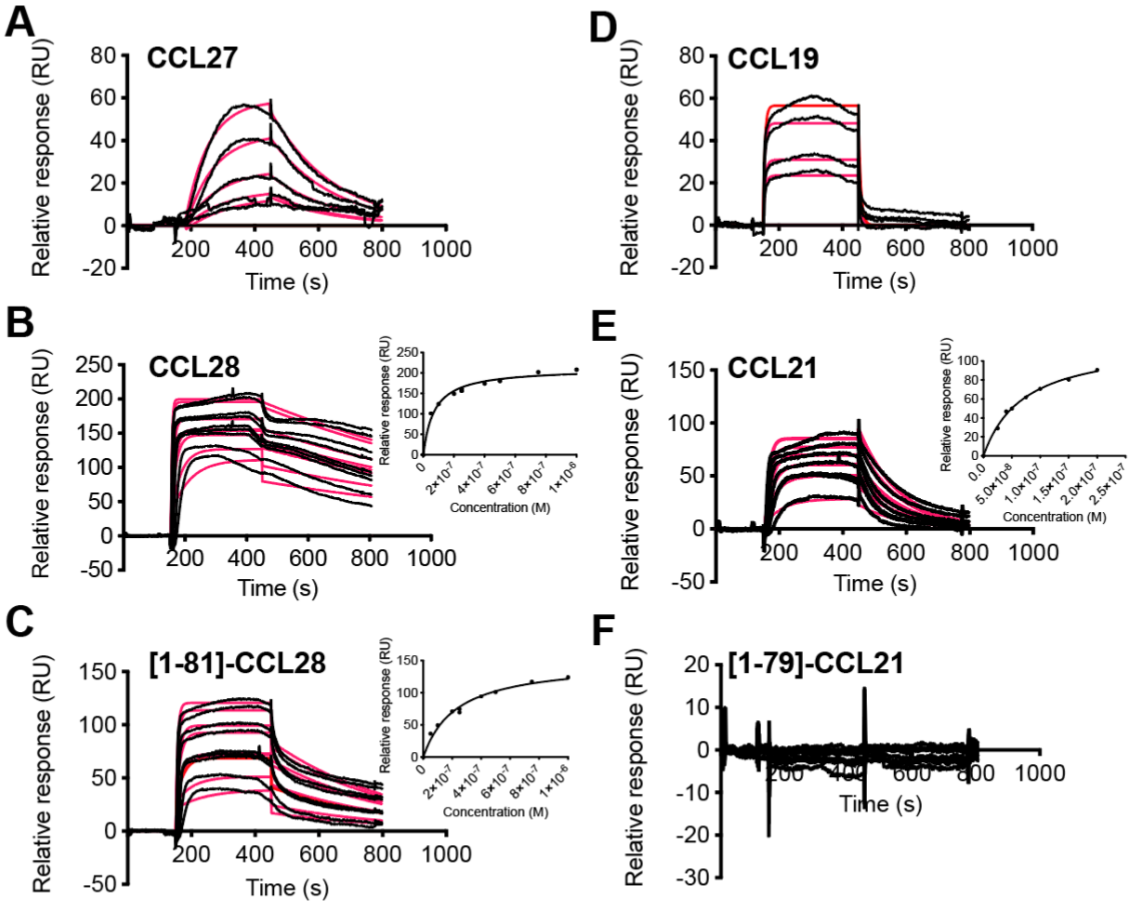
